# Autophagy disruption and mitochondrial stress precede photoreceptor necroptosis in multiple mouse models of inherited retinal disorders

**DOI:** 10.1101/2024.08.02.606303

**Authors:** Fay Newton, Mihail Halachev, Linda Nguyen, Lisa McKie, Pleasantine Mill, Roly Megaw

## Abstract

Inherited retinal diseases (IRDs) are a leading cause of blindness worldwide. One of the greatest barriers to developing treatments for IRDs is the heterogeneity of these disorders, with causative mutations identified in over 280 genes. It is therefore a priority to find therapies applicable to a broad range of genetic causes. To do so requires a greater understanding of the common or overlapping molecular pathways that lead to photoreceptor death in IRDs and the molecular processes through which they converge. Here, we characterise the contribution of different cell death mechanisms to photoreceptor degeneration and loss throughout disease progression in humanised mouse models of IRDs. Using single-cell transcriptomics, we identify common transcriptional signatures in degenerating photoreceptors. Further, we show that in genetically and functionally distinct IRD models, common early defects in autophagy and mitochondrial damage exist, triggering photoreceptor cell death by necroptosis in later disease stages. These results suggest that, regardless of the underlying genetic cause, these pathways likely contribute to cell death in IRDs. These insights provide potential therapeutic targets for novel, gene-agnostic treatments for IRDs applicable to the majority of patients.

## Introduction

Inherited retinal disorders (IRDs) are the most common cause of blindness in children and adults of working age, affecting 1 in 1380 people,(1) and characterised by death of our light-sensing photoreceptors. Retinitis pigmentosa (RP) is the most common form, resulting in an initial loss of rod photoreceptors (followed by non-autonomous cone photoreceptor death)(2, 3), whilst other IRDs, such as cone dystrophies, cone-rod dystrophies and macular dystrophies, preferentially affect cones first. Although the clinical features of IRDs are well characterised, it is a highly genetically heterogeneous disorder with causative mutations identified in more than 280 genes(4). Therefore, despite recent advances in gene replacement and genome editing therapies for IRDs (5, 6), developing treatments that target common downstream disease mechanisms, and are thus applicable to a broader range of causative alleles in a gene-agnostic manner, is a strategic priority for eye health for all IRD patients.

IRDs are caused by mutations in genes encoding a diverse range of proteins, with roles in phototransduction, rhodopsin cycling, photoreceptor structure and intracellular trafficking(7–9). Whilst our knowledge of the function of these genes has improved, there is comparatively little understanding of how disruption of their function impacts on cellular processes that in turn lead to photoreceptor death(10). Dysregulation of autophagy, the lysosome-dependent pathway through which damaged cell contents are degraded, has been implicated in some IRD models(11–13). Further, some studies have identified apoptosis as the major cell death pathway in IRDs(14–16), while others demonstrate that alternate forms of regulated cell death such as necroptosis, ferroptosis and parthanatos play significant roles(17–22). In order to protect photoreceptors in IRDs using drug-based strategies, it will be vital to define the common mechanisms of photoreceptor death and the upstream pathways from which they are triggered.

Here, using single-cell RNA sequencing (scRNAseq), RP mouse models and multiple imaging modalities, we provide evidence supporting a model whereby photoreceptors are subject to increased mitochondrial stress early in disease. Further, we show that this leads to defects in autophagy. Finally, we demonstrate that photoreceptors undergo programmed cell death via the necroptosis pathway and that similar pathways could contribute to cell death in RP, regardless of the underlying genetic cause. These insights provide potential therapeutic targets to treat RP in patients carrying a wide range of causative alleles.

### Single-cell transcriptomes reveal an *Rpgr* mutant-specific population of macrophages and distinct sub-populations of rod photoreceptors

RPGR functions to regulate outer segment disc turnover, with pathogenic mutations causing a spectrum of retinal disease, from RP to cone-rod dystrophy to cone dystrophy. In keeping, we previously generated two novel *Rpgr* mutant mouse lines that replicate human pathogenic mutations and exhibit similar disease kinetics to human patients(23). *Rpgr*^*Ex*3*d*8^, which harbours an 8 base pair mutation in exon 3 (resulting in a premature termination codon in all transcripts, thus is a null allele with no expression), initially manifests cone disease. *Rpgr*^*ORF d*5^, which has a 5 base pair truncation mutation within the C-terminal repetitive domain of the retina-specific isoform, initially manifests rod disease. Both lines exhibit retinal stress, indicated by increased GFAP upregulation in Müller glia (**Supplementary Fig. 1**), prior to significant photoreceptor degeneration at 18 months(23). These lines thus offer excellent models with which to identify shared pathway changes across the IRD spectrum.

To profile different retinal cell states that occur as photoreceptors degenerate, we performed scRNAseq on retinas of 18-month-old *Rpgr*^*ORF d*5^ and *Rpgr*^*Ex*3*d*8^ mice and wildtype littermates. Cluster assignment using Seurat identified clusters corresponding to the major retinal cell types (**Supplementary Fig. 2a, 3a**), classified by expression of known markers of each cell type (**Supplementary Table 1, Supplementary Fig. 2c, 3c**). Comparison of the expression profiles of mutant and wild-type cells in rod photoreceptor clusters showed that genes required for photoreceptor function were downregulated in mutant cells compared to wild type (**Supplementary Fig. 2d, 3d**), with retinal degenerative diseases being the most highly enriched GO term (**Supplementary Fig. 4b**).

In addition, analysis of the *Rpgr*^*ORF d*5^ scRNAseq data identified a small cluster, characterised by expression of macrophage markers (**Supplementary Fig. 2a, b**), exclusively composed of cells from the mutant sample. Immunofluorescent staining of retina cryosections confirmed cells, positive for the pan-macrophage marker F4/80 and the microglial marker TMEM119, invading outer layers of *Rpgr*^*ORF d*5^ and *Rpgr*^*Ex*3*d*8^ retinas at 18 months, something not observed in wild type at this age (**Supplementary Fig. 2e, 3e**). Whilst double positive cells are likely nonmigratory, tissue-resident macrophages, a subset of F4/80+; TMEM119cells are seen, suggesting that circulating monocyte-derived macrophages also contribute to this cell cluster. It is not clear why this population was not present in the *Rpgr*^*Ex*3*d*8^ scRNAseq data. However, since immunofluorescence shows there to be few F4/80-positive cells in the outer retina, the profile may have been lost during quality control stages.

Interestingly, both scRNAseq experiments identified several sub-clusters of rod photoreceptors (**Fig 1a, Supplementary Fig. 2a, 3a**). Whilst cells in all sub-clusters expressed rod photoreceptor markers (**Supplementary Fig. 2c, 3c**), each sub-cluster showed differential gene expression, suggesting distinct expression profiles. Similar characteristic genes for each of these sub-clusters were identified in both scRNAseq experiments (**Supplementary Table 1**). Pseudotime analysis using Slingshot showed that these sub-clusters follow a trajectory whereby expression of genes required for phototransduction decreases between adjacent clusters (**Fig. 1a, b**). These sub-clusters may therefore represent rod photoreceptors performing sub-optimally, at different stages of degeneration. Indeed, in the *Rpgr*^*ORF d*5^ experiment, clusters with reduced phototransduction gene expression contained a higher percentage of photoreceptors from the mutant sample (**Supplementary Table 2, Supplementary Fig. 2b**). The presence of wild-type cells in these clusters could indicate age-related functional decline.

**Fig. 1.**
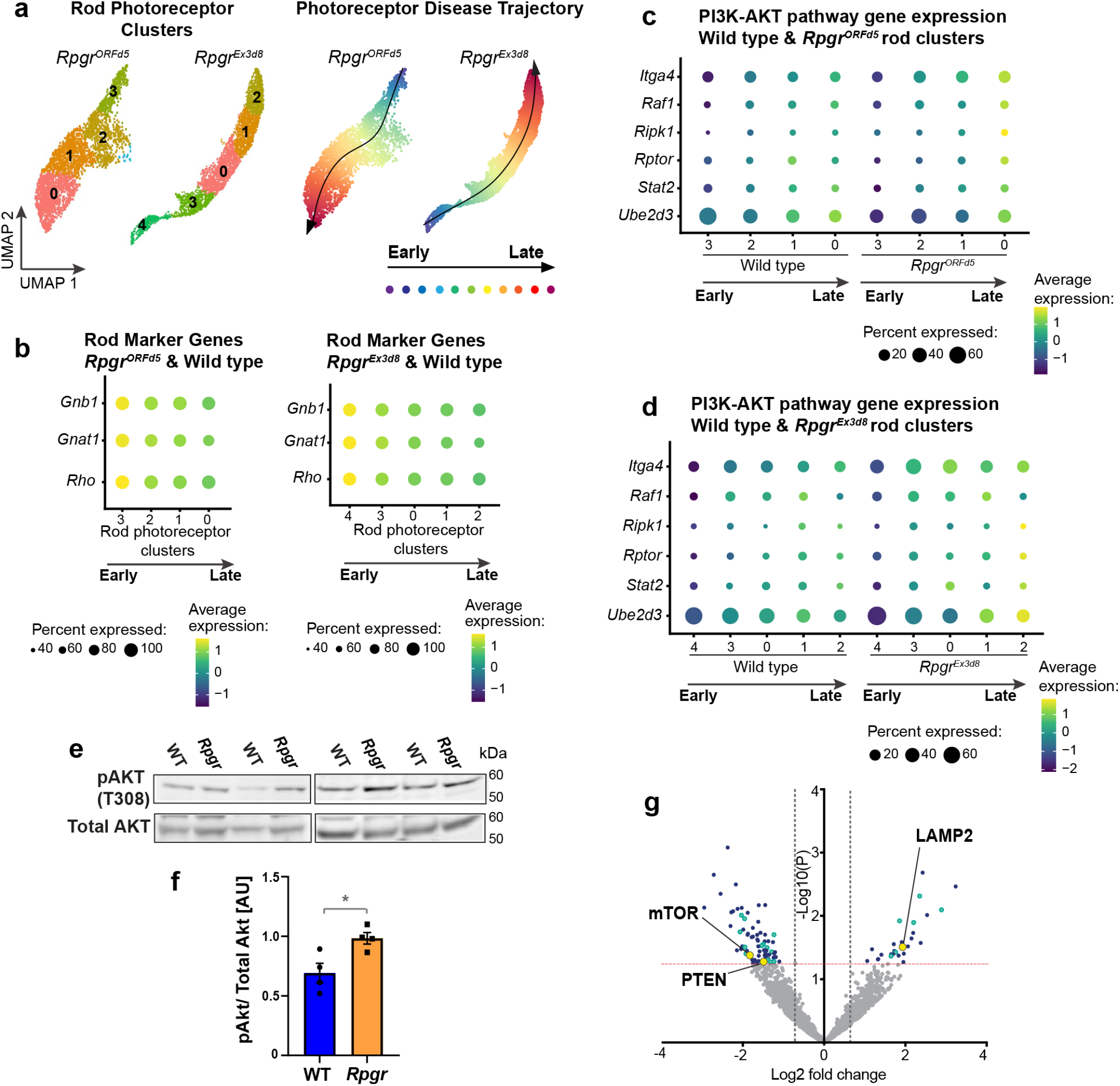
Single-cell transcriptomics identifies photoreceptor populations with reduced phototransduction gene expression and increased PI3K/AKT pathway gene expression. **a** Subclusters of rod photoreceptors numbered by Seurat according to cluster size (number of cells in each cluster; left panel). Pseudotime analysis of these clusters shows photoreceptor disease trajectory (right panel). **b** Genes required for phototransduction are downregulated along this trajectory (combined mutant and wild type cells). **c, d** Genes associated with PI3K/AKT signalling are upregulated in late (more degenerated) clusters compared to early (less degenerated) clusters. This downregulation is more pronounced in *Rpgr*^*ORF d*5^ **(c)** and *Rpgr*^*Ex*3*d*8^ **(d)** mutant photoreceptors compared to wild type. **e, f** Active pAKT is increased in *Rpgr*^*Ex*3*d*8^ mutants relative to total AKT (Bars show mean; N = 4 mice per experimental group; error bars show SEM; * p = 0.037). **g** Mass spectrometry analysis of *Rpgr*^*Ex*3*d*8^ retina lysate compared to WT at 6 months (data from Megaw et al., 2024). Red dotted line indicates p < 0.05, grey dotted lines indicate Log2 fold change < -0.6 and > 0.6. PTEN, mTOR and LAMP2 highlighted by yellow circles, proteins associated with cell stress pathways are highlighted in cyan (listed in Supplementary table 4). Other proteins with significant fold change (Log2 fold change ≤ −0.6*or* ≥ 0.6,p ≤0.05) are highlighted in blue.

Next, we analysed genes that showed increased photoreceptor expression along this disease trajectory. KEGG pathway analysis revealed up-regulation of mRNA processing, TNF-α/NF-κB signalling and lysosome biogenesis pathways (**Supplementary Fig. 4a**) suggesting increased activation of inflammatory pathways and increased requirement for lysosomes, which may indicate increased or dysregulated autophagy. Interestingly, we also found several genes with increased expression in more degenerated clusters have a role in PI3K/AKT signalling (**Supplementary Fig. 4c, d**), a pathway associated with cell survival and regulation of autophagy. Expression of these genes increased progressively in wild-type photoreceptors from ‘early’ to ‘late’ clusters but this upregulation was more pronounced in mutant photoreceptors (**Fig. 1c, d**). To further interrogate this pathway, we looked specifically in *Rpgr*^*Ex*3*d*8^ retina, which develops a faster retinal degeneration than *Rpgr*^*ORF d*5^. We observed an increase in active, phosphorylated AKT (pAKT) relative to total AKT in retinal lysates from *Rpgr*^*Ex*3*d*8^ mutants compared to wild type at 12 months (**Fig. 1e, f**). Further, mass spectrometry analysis of total retinal lysates from six-monthold *Rpgr*^*Ex*3*d*8^ mice 23 showed decreased expression of the PI3K/AKT pathway inhibitor PTEN and the downstream effector mTOR, compared to wild-type littermates (**Fig. 1g**). We therefore sought to characterise dysregulation in pathways governed by PI3K/AKT.

### *Rpgr* mutant photoreceptors accumulate autophagosomes

The PI3K/AKT pathway regulates autophagy, the lysosome-dependent pathway through which damaged cell contents are degraded, which has been implicated in photoreceptor degeneration in some RP models(11–13). In addition to the dysregulation of PI3K/AKT and lysosomal biogenesis pathways, our scRNAseq data also revealed changes in expression of autophagy pathway genes in *Rpgr*^*Ex*3*d*8^ and *Rpgr*^*ORF d*5^ mutant photoreceptors compared to wild type (**Fig. 2a, b**). To further explore autophagy regulation in RP, we focused further functional experiments on the faster degenerating *Rpgr*^*Ex*3*d*8^ model. We performed transmission electron microscopy (TEM), which revealed a significant increase in large, vesicle-like structures in mutant photoreceptor inner segments compared to wild-type controls at 6 and 12 months of age (**Fig. 2c, d**). This suggested an accumulation of autophagosomes or autolysosomes, indicating a defect in autophagy, and was supported by elevated autophagy marker p62 (also known as SQSTM1) in photoreceptor inner segments at 12 months in our *Rpgr*^*Ex*3*d*8^ mutant (**Fig. 3a, b**). Interestingly, this accumulation was not present at 18 months (**Fig. 3b**). Similarly, there was significant accumulation of the lysosomal protein LAMP1 in *Rpgr*^*Ex*3*d*8^ photoreceptor inner segments at 12 months, but not at 18 months (**Fig. 3c, d**). In further support of this, another lysosomal protein, LAMP2, was significantly upregulated in mutant retinas in our total proteomic data (**Fig. 1g**). We conclude, therefore, that there is increased autophagy in *Rpgr* mutant photoreceptors in mice up to 12 months of age.

**Fig. 2.**
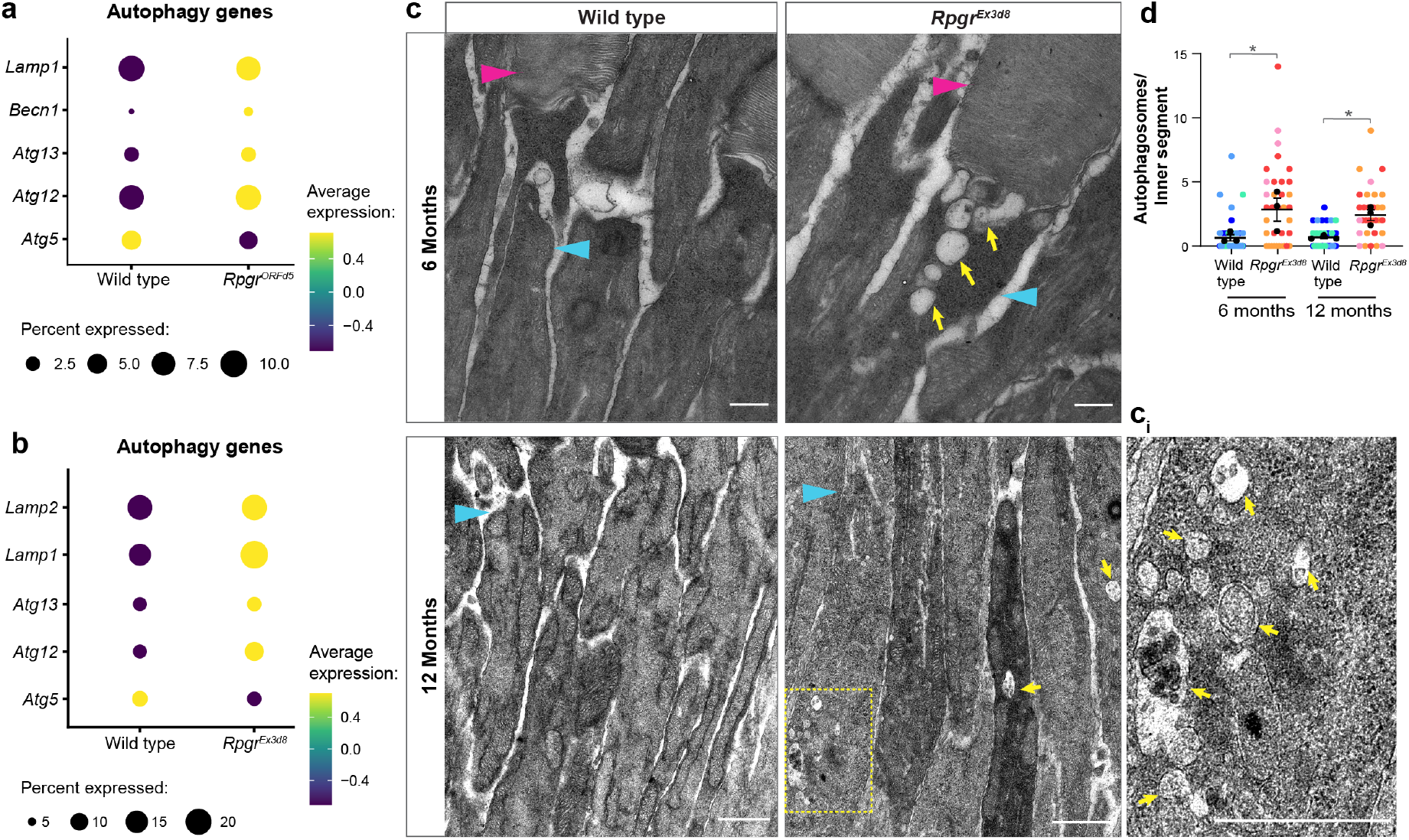
*Rpgr* mutant photoreceptors accumulate autophagosomes in inner segments. **a, b** Changes in expression level of autophagy genes in *Rpgr*^*ORF d*5^ **(a)** and *Rpgr*^*Ex*3*d*8^ mutant **(b)** rod photoreceptors compared to wild type in scRNAseq data. **c** Transmission electron microscopy images of *Rpgr*^*Ex*3*d*8^ and wild-type littermate control photoreceptor inner segments at 6 and 12 months of age show accumulation of autophagosomes in *Rpgr*^*Ex*3*d*8^ mutant photoreceptors (magenta arrowheads = outer segments, cyan arrowheads = inner segments, yellow arrows and yellow boxed region indicate autophagosomes, yellow boxed region enlarged in ci, Scale bar = 500 nm.) **d** Quantification of autophagosome numbers per inner segment. Colours indicate autophagosome counts per photoreceptor from individual mice, with mean values for each mouse superimposed in black (N = 3 animals per genotype; n > 5 measurements per animal) error bars show SEM; * p = 0.039 (6 months), p = 0.041 (12 months)).

**Fig. 3.**
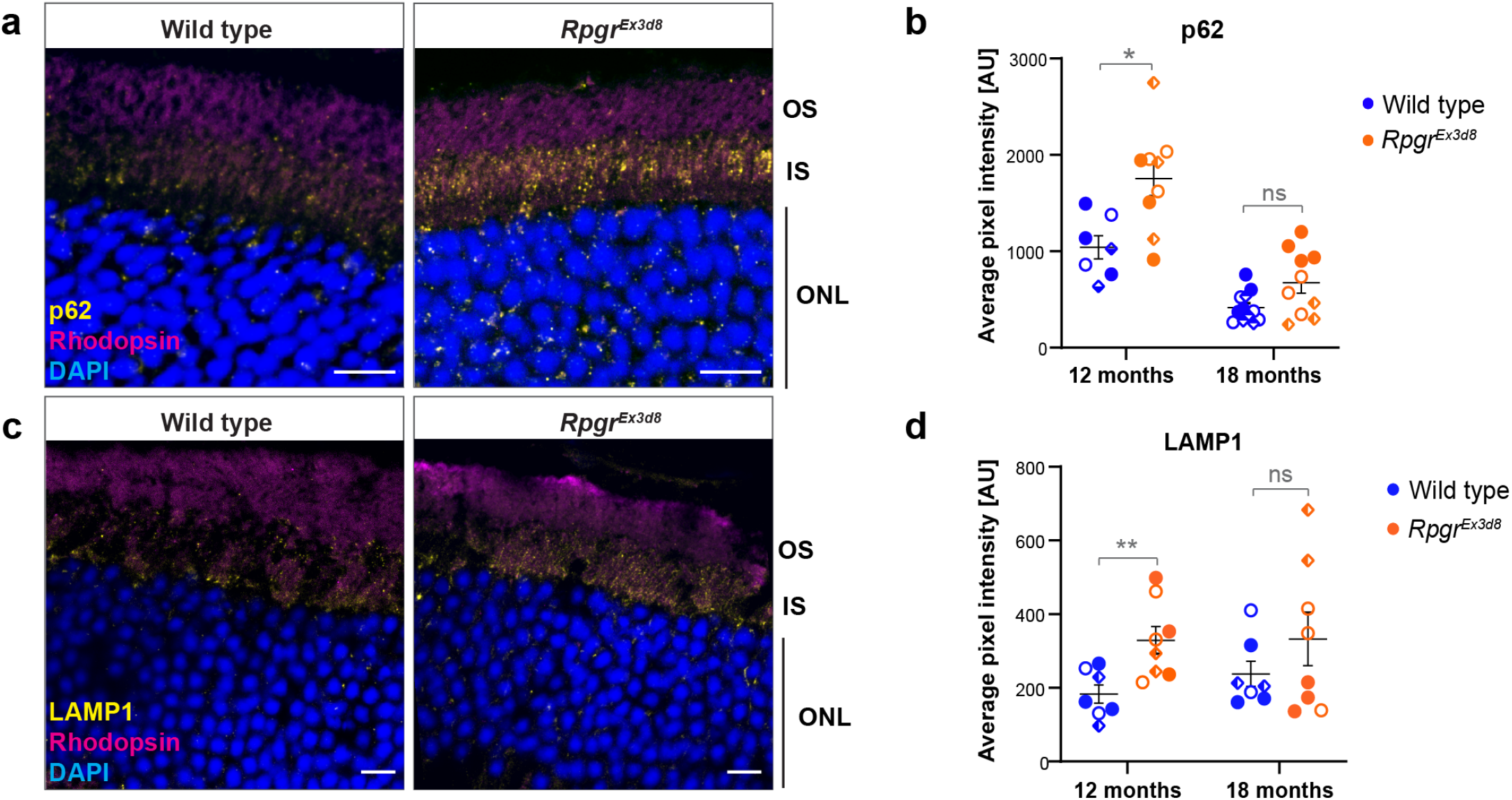
Autophagosome and lysosome markers accumulate in *Rpgr* mutant photoreceptor inner segments. **a** Accumulation of p62 in *Rpgr*^*Ex*3*d*8^ photoreceptor inner segments (scale bar = 10 µm). **b** Quantification of p62 staining in inner segment. **c** Accumulation of LAMP1 in *Rpgr*^*Ex*3*d*8^ photoreceptor inner segments (scale bar = 10 µm). **d** Quantification of LAMP1 staining in inner segment. (b, d; symbols indicate pixel intensity per image from individual mice (N = 3 animals per genotype; n > 2 sections per animal (symbol) error bars show SEM; * p = 0.042, ** p = 0.007)).

### *Rpgr* mutant photoreceptors exhibit mitochondrial damage

To gain further insight into cellular changes occurring at later stages of degeneration, we carried out TEM on retinas from *Rpgr*^*Ex*3*d*8^ mice at 12 and 18 months, comparing them to wild-type littermates. At 12 months, when accumulation of autophagosomes was most prevalent in mutant photoreceptors (**Fig. 2c**), we also observed occasional mitochondrial defects such as swelling or membrane blebbing (**Fig. 4a**). At 18 months, by contrast, when autophagosome numbers were similar to that of wild type, severe swelling of *Rpgr*^*Ex*3*d*8^ mitochondria was observed, indicative of mitochondrial stress in mutant photoreceptors (**Fig. 4b**). Further, mitochondrial genes were up-regulated in both *Rpgr*^*ORF d*5^ and *Rpgr*^*Ex*3*d*8^ mutant photoreceptors in our scRNAseq data (**Fig. 4c, d**). One of these genes, Vdac1, encodes a mitochondrial outer membrane (MOM) calcium channel protein with a role in the mitochondrial stress response(24). VDAC1 forms part of the MOM complex required for crosstalk with the endoplasmic reticulum (ER) and regulates apoptosis, autophagy and mitophagy(25, 26). Under stress conditions, VDAC1 is up-regulated and forms oligomers at the MOM, increasing membrane permeability and promoting cytochrome C release(27, 28). Immunofluorescent staining showed increased levels of VDAC1 in *Rpgr*^*Ex*3*d*8^ mutant photoreceptor inner segments at 12 and 18 months, supporting increased mitochondrial stress (**Fig. 5a-c**).

**Fig. 4.**
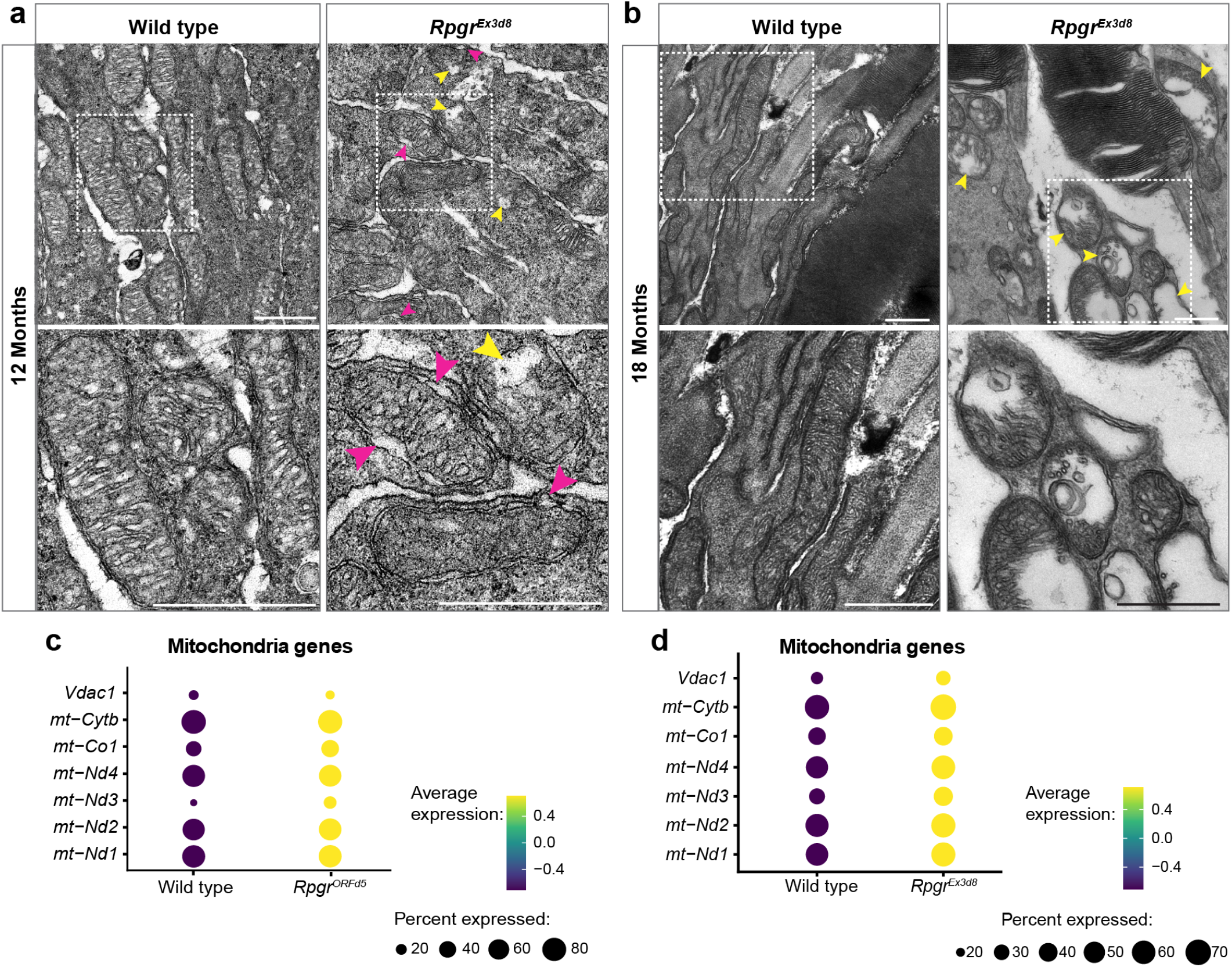
Rpgr-mutant photoreceptors have abnormal mitochondrial morphology, indicative of mitochondrial damage. Transmission electron microscopy images of mutant and wild-type photoreceptor inner segments at 12 months **(a)** and 18 months **(b)** show mitochondrial swelling (yellow arrowheads) and blebbing of mitochondrial membranes (magenta arrowheads) at 12 months in mutant retina. Boxed regions enlarged in lower panels (scale bar = 500 nm). **c, d** Increased expression of genes required for mitochondrial function in *Rpgr*^*ORF d*5^ **(c)** and *Rpgr*^*Ex*3*d*8^ **(d)** mutant photoreceptors in scRNAseq data.

**Fig. 5.**
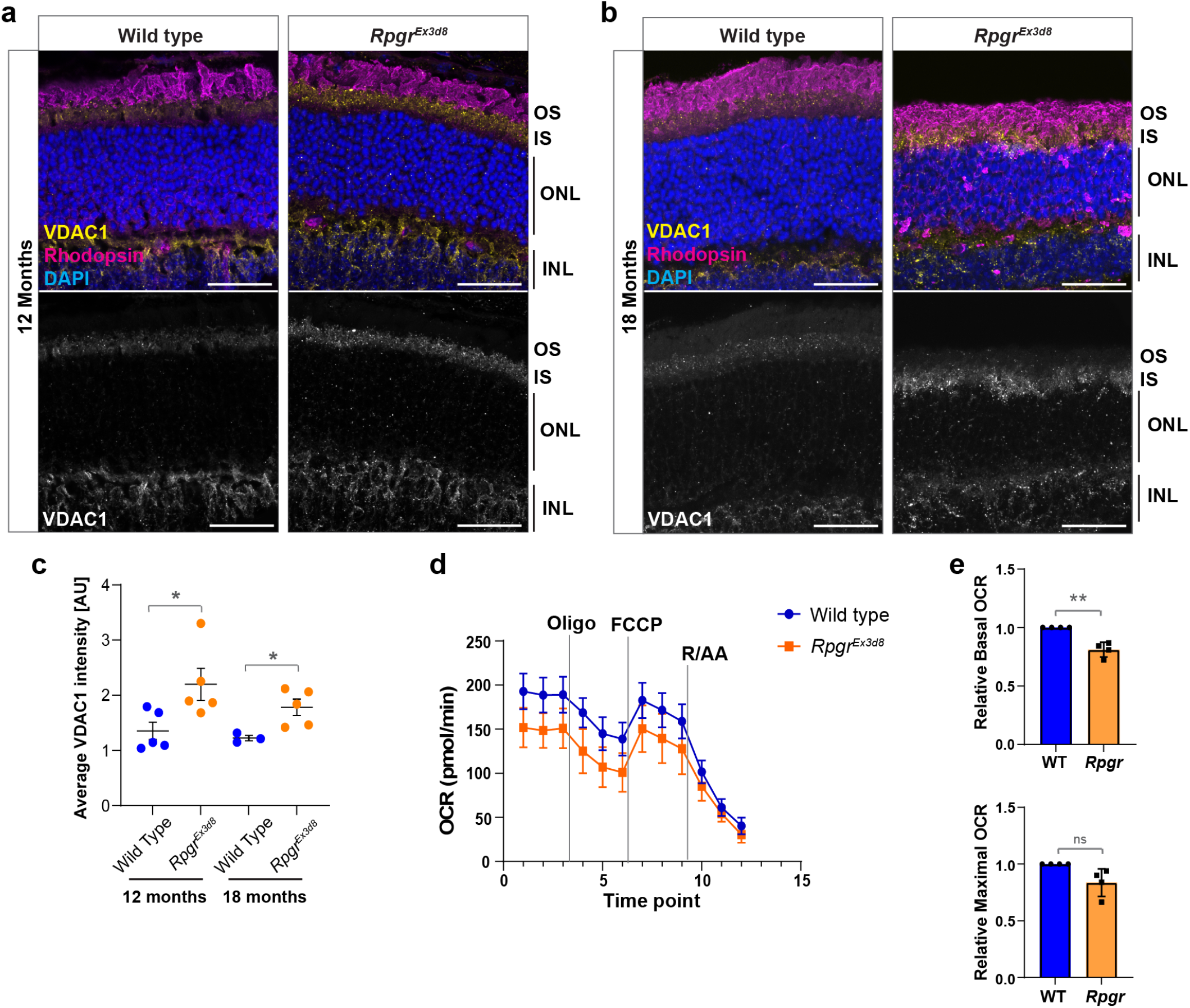
Mitochondrial stress develops in Rpgr mutant photoreceptors. **a, b** VDAC1 expression is increased in mutant photoreceptor inner segments at 12 **(a)** and 18 **(b)** months compared to wild-type littermate controls (scale bar = 25 µm.) **c** Quantification of VDAC1 staining at 12 and 18 months (N = 3 animals per genotype; error bars show SEM; * p = 0.042 (12 months), p = 0.017 (18 months)) **d** Mitochondrial stress assay shows basal oxygen consumption rate (OCR) is reduced in *Rpgr*^*Ex*3*d*8^ mutants. Mean OCR for 4 biological replicates (N = 4 animals per genotype; 10 biopsies per animal) is shown at each time point (error bars show SD). Vertical lines indicate points of addition of relevant modulators (oligo = oligomycin; FCCP = carbonyl cyanide p-trifluoro-methoxyphenyl hydrazone; R/AA = rotenone + antimycin A). **e** Mean OCR of mutant retina (time point 3 = Basal, time point 7 = Max) normalised to the mean OCR of wild-type littermate controls at the same time point (error bars show SD; ** p = 0.009).

To directly assess mitochondrial respiration in *Rpgr* mutants, we measured oxygen consumption rate (OCR) in ex-vivo retinal punches from 14-month-old *Rpgr*^*Ex*3*d*8^ mice and wildtype littermates. We observed a significant reduction in basal OCR in *Rpgr*^*Ex*3*d*8^ retinas, suggesting reduced mitochondrial function (**Fig. 5d, e**). We found no difference between basal and maximal OCR in either mutant or wild-type retinas after adding an uncoupler (FCCP) and therefore could not measure mitochondrial reserve capacity. However, this is consistent with previous studies showing that, due to high oxygen demand, photoreceptor mitochondria function close to maximum capacity(29). Taken together, these findings support that *Rpgr*^*Ex*3*d*8^ mutant photoreceptors develop mitochondrial stress from 12 months, worsening as the retina degenerates.

### Necroptosis is the major cell death pathway in *Rpgr* photoreceptors

Mitochondrial stress is associated with increased apoptosis, due to release of cytochrome C from mitochondria(27). To determine whether apoptosis contributes to the high level of photoreceptor death in *Rpgr* mutants at later stages, 18-month retinal sections were stained with antibodies for cleaved caspase 3 (cl-casp3). We observed increased clcasp3 positive photoreceptors in mutant retinas compared to wild type (**Supplementary Fig. 5a**). However, the relatively small number of positive cells seemed insufficient to explain the significant loss of photoreceptors observed at this age 23 and analysis of our scRNAseq revealed no obvious apoptotic transcriptional signature (**Supplementary Fig. 5b, c**). This suggested that apoptosis may not be the primary mechanism of photoreceptor death in *Rpgr* mutants. In support of this, *Rpgr*^*Ex*3*d*8^ mutant retinas have reduced cleaved caspase 8 (cl-casp8) relative to the full-length protein (fl-casp8, (**Supplementary Fig. 5d, e**), suggesting reduced activation of apoptosis. An alternative cell death pathway was sought.

Gene ontology differential expression analysis of our scRNAseq data suggested that both *Rpgr*^*ORF d*5^ and *Rpgr*^*Ex*3*d*8^ mutant photoreceptors had upregulation of necroptosis-associated genes (**Fig. 6a, b, Supplementary Fig 4c**), a form of regulated cell death characterised by high inflammation and cell membrane rupture. Necroptosis can also be induced by mitochondrial stress(30). We therefore investigated whether necroptosis could be involved in photoreceptor death in *Rpgr* mutants. Immunofluorescence of pMLKL, the terminal protein in the necroptotic cell death programme, revealed increased pMLKL positive photoreceptors in *Rpgr*^*Ex*3*d*8^ and *Rpgr*^*ORF d*5^ mutants, with pMLKL accumulating at the membrane around the cell body (**Fig. 6c, d, Supplementary Fig 6**). A much larger number of photoreceptors were affected compared to the number of photoreceptors positive for cl-casp3 at 18 months (**Fig. 6d, Supplementary Fig 6b**), suggesting necroptosis is the major cell death pathway in *Rpgr* mutant models of RP. The number of pMLKL positive cells also increased progressively from 6-18 months (**Fig. 6d, Supplementary Fig 6b**), corresponding to the gradual loss of photoreceptors over this time period.

**Fig. 6.**
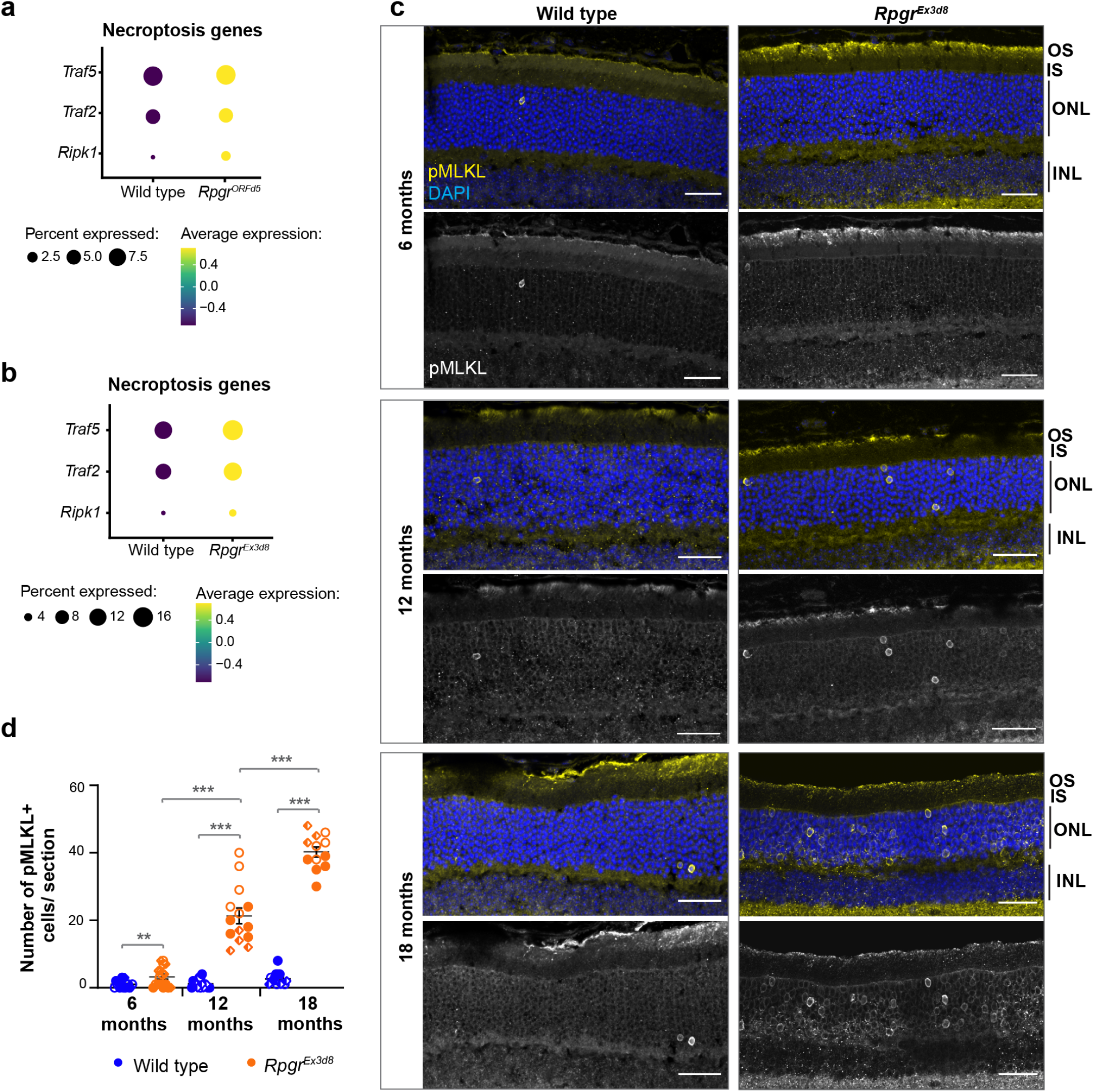
Photoreceptors die by necroptosis in Rpgr mutants. **a, b** Necroptosis genes have increased expression in *Rpgr*^*ORF d*5^ **(a)** and *Rpgr*^*Ex*3*d*8^ **(b)** mutant rod photoreceptors in scRNAseq. **c** Increased pMLKL positive photoreceptors in *Rpgr*^*Ex*3*d*8^ mutants at 6, 12 and 18 months compared to wild-type littermate controls (all scale bars = 25 µm). **d** Quantification of pMLKL positive photoreceptors (symbols indicate images from individual mice (N = 3 animals per genotype; n > 4 sections per animal (symbol); bars show mean; error bars show SEM; ** p = 0.009; *** p = 7.85E-7 (12 months), p = 1.74E-12 (18 months), p = 1.78E-6 (6 vs 12 months), p = 7.63E-7 (12 vs 18 months)).

To test whether necroptosis is also an important mechanism in other RP models, we examined pMLKL expression in the *Pde*6*b*^*atrd*2^ mouse(31). In contrast to RPGR’s role in photoreceptor outer segment maintenance, the PDE6 protein complex is involved in the phototransduction cascade and thus represents a suitably distinct model with which to probe for gene-agnostic, shared pathway disruptions.*Pde6b* mutations result in much faster photoreceptor degeneration, with the *Pde*6*b*^*atrd*2^ mouse showing extreme thinning of the ONL by P28(31). pMLKL expression was significantly higher in *Pde*6*b*^*atrd*2^ photoreceptors and increased progressively as they die, from P14 to P28 (**Fig. 7a, b**). Further, *Pde*6*b*^*atrd*2^ photoreceptors showed progressive defects in mitochondrial morphology from P13-P18, similar to those observed in older *Rpgr* mutants (**Fig. 7c**). We also observed increased prevalence of enlarged autophagosomes at P13 (**Fig. 7c**), as well as accumulation of p62 in *Pde*6*b*^*atrd*2^ photoreceptor inner segments at P14 and P16 (**Fig. 7d, e**). We conclude, therefore, that dysregulation of autophagy, mitochondrial stress and necroptosis are the key mechanisms contributing to photoreceptor death in multiple RP models, regardless of the underlying cause and gene function (**Supplementary Fig. 7**).

**Fig. 7.**
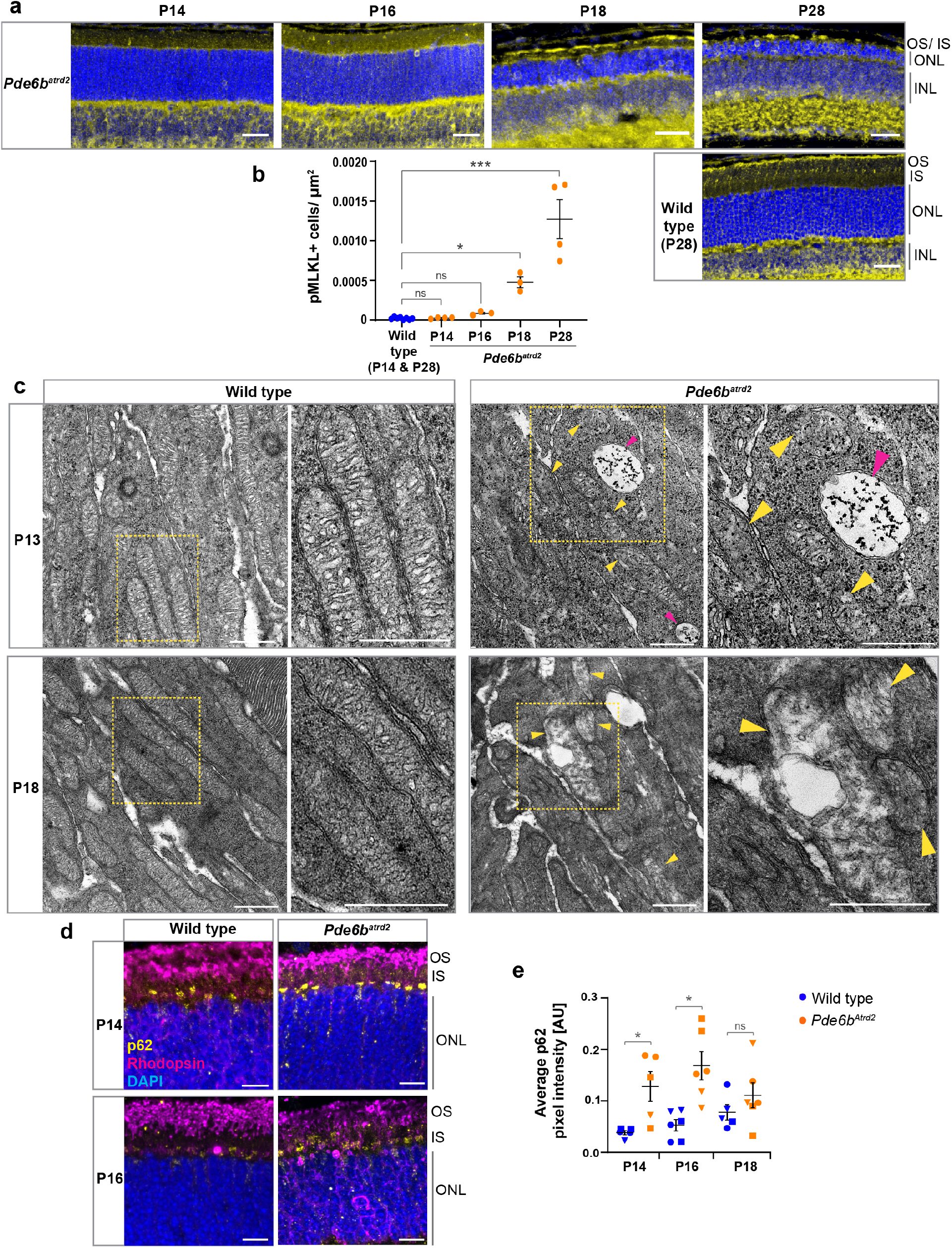
Figure 7: Necroptosis, mitochondrial stress and autophagy dysregulation are important cell death mechanisms in the *P de*6*b*^*atrd*2^ mouse model of RP. **a** Increased numbers of pMLKL positive photoreceptors were observed at postnatal day 14, 16, 18 and 28 (scale bar = 25 µm). **b** Quantification of pMLKL positive photoreceptors per µm2 in each retinal section. Wild type combines counts from P14 and P28 retinas from different mice (N = 8 animals in total). All other data points are from individual mice (P14 and P28: N = 4 animals per time point; P16 and P18: N = 3 animals per time point; bars show mean, error bars show SEM; * p = 0.027; *** p = 0.0006. **c** Transmission electron microscopy images of mutant and wild-type photoreceptor inner segments show mitochondrial swelling in *P de*6*b*^*atrd*2^ mutant photoreceptors (yellow arrowheads) at P13 and P18. Large autophagosomes are also present in *P de*6*b*^*atrd*2^ photoreceptor inner segments at P13 (magenta arrowheads) (yellow boxed regions enlarged in right hand panels; scale bars = 500 nm). **d** Accumulation of p62 in *P de*6*b*^*atrd*2^ photoreceptor inner segments (scale bar = 10 µm). **e** Quantification of p62 staining in inner segment. (Symbols indicate pixel intensity per image from individual mice; N = 3 animals per genotype at each time point; error bars show SEM; * p = 0.036 (P14), p = 0.016 (P16)).

## Discussion

Understanding the processes underlying photoreceptor degeneration in RP remains an important challenge for the development of effective therapies. Here, we identify mitochondrial stress and dysregulated autophagy as major contributors to photoreceptor deterioration in *Rpgr* and *Pde*6*b*^*atrd*2^ mutants, with most cells eventually dying by necroptosis (**Supplementary Fig. 7**). In contrast to apoptosis, necroptosis is a form of regulated necrosis involving cellular membrane rupture, release of damage associated molecular patterns (DAMPs) and, consequently, a proinflammatory environment resulting in damage to surrounding tissues(32, 33). This is in keeping with the clinical picture of RP, where a chronic, low-grade inflammation often leads to posterior, sub-capsular cataract, epiretinal membrane formation and cystoid macular oedema.

We found that PI3K/AKT signalling is upregulated in *Rpgr* mutant photoreceptors. PI3K/AKT signalling is generally associated with cell survival; activation of AKT inhibits apoptosis(34–36) and could therefore represent a protective mechanism in response to cell stress. However, sustained activation of AKT triggers necroptosis in neuronal cells(37) and could play a similar role in *Rpgr* mutant photoreceptors. Sev-eral other studies, using different models of RP, have identified necroptosis as the primary mechanism of photoreceptor death(17, 18, 20). Our finding, therefore, that necroptosis has a significant role in photoreceptor loss in multiple RP mouse models supports that this mechanism may be important in RP, regardless of the genetic cause, and its inhibition could be an attractive target for development of future therapies.

mTOR, reduced in our mutant retinas, acts to inhibit autophagy(36, 38). Depletion of mTOR, therefore, could explain the increased autophagy observed in *Rpgr* mutant photoreceptors at early disease stages. We also observed increased autophagy in *P de*6*b*^*atrd*2^ photoreceptors at early disease stage, in keeping with other *Pde6b* models(39), suggesting dysregulation of autophagy could play an important role in early photoreceptor degeneration arising from differ-ent genetic causes. This increase in autophagy may be a protective response to remove misfolded or mis-localised proteins such as rhodopsin (**Supplementary Fig. 1**). It could also be an attempt to remove the damaged mitochondria seen in our mouse models, which would be in keeping with the increased oxidative stress seen in our mutant scRNAseq and mass spectrometry datasets. Indeed, metabolic and oxidative stress appear to play significant roles in photoreceptor degeneration in several other models of RP(40–44). Further work is needed to resolve the content of the observed autophagosomes. However, photoreceptor autophagy is complex, and inhibition of autophagy appears protective in certain mouse RP models(12).

Alternatively, the autophagosome and lysosome accumulation we observed could indicate progressively defective autophagy, with stalled autophagosomes failing to clear stressed mitochondria and triggering cell death. This could explain why significant accumulation of p62 and LAMP1 was not observed at 18 months, when severely affected photoreceptors have been lost. Photoreceptors have a high basal level of autophagy(45) and may therefore be particularly sensitive to disruption of the dynamic process of autophagy, termed autophagic flux. Further work is needed to define the state of autophagy flux in RP.

While mitochondrial dysfunction is not required to initiate necroptosis(46), mitochondrial reactive oxygen species (ROS) have been shown to promote autophosphorylation of receptor interacting protein kinase 1 (RIPK1) and can therefore facilitate formation of the RIPK1/RIPK3/pMLKL necrosome complex(47, 48). RIPK3 can subsequently activate the mitochondrial pyruvate dehydrogenase complex, leading to enhanced aerobic respiration and increased generation of ROS(49, 50). This acts as a feedforward mechanism, further increasing necroptosis. Mitochondrial stress in *Rpgr* mutants could therefore facilitate photoreceptor death by necroptosis. Indeed, inhibition of the mitochondrial stress response was shown to alleviate necroptosis in rat retinal neurons following ischemia-reperfusion injury(30). The autophagy machinery can also form a scaffold for the necrosome, mediated by p62 recruitment of RIPK1, and this interaction promotes cell death by necroptosis rather than apoptosis(51). It is therefore possible that the need to remove damaged mitochondria in *Rpgr* mutant photoreceptors leads to enhanced mitophagy, which in turn promotes necroptosis. Interestingly, an increased association of the autophagy regulators BECN1 and ATG5 with mitochondria has been shown in photoreceptor inner segments in a RhoP23H rat model of RP, suggesting increased mitophagy. Necroptosis was also identified as the primary mechanism of photoreceptor death in this model(20). Necroptosis has a role in other neurodegenerative diseases, where autophagy dysregulation contributes to the pathology(52–56), and inhibition of necroptosis can reduce neural degeneration in animal models(52, 54, 56–59) and xenografted human neurons(60). A greater understanding of the interplay between necroptosis, autophagy/mitophagy and mitochondrial stress in RP, therefore, will allow identification of potential therapeutic targets upstream of necroptosis pathway activation.

## Methods

### Animals

*Rpgr*^*Ex*3*d*8^ and *Rpgr*^*ORF d*5^ mice were generated using CRISPR/Cas9 genome editing as described in Megaw et al. 2024. *Pde*6*b*^*atrd*2^ mice were generated by N-ethyl-Nnitrosourea (ENU) induced mutation as described (31). All experiments followed international, national and institutional guidelines for the care and use of animals. Animal experiments were carried out under UK Home Office Project Licenses PPL P1914806F and PP5301383 in facilities at the University of Edinburgh (PEL 719 60/2605) and were approved by the University of Edinburgh animal welfare and ethical review body.

### Single-cell RNA sequencing and Analysis

Mice were sacrificed, eyes enucleated and placed into PBS. Retinas were dissected in NeuroCult Tissue Collection Solution (Stem Cell Technologies) and dissociated into single cells using the NeuroCult Enzymatic Dissociation Kit for Adult CNS Tissue (Stem Cell Technologies cat. 05715) according to the manufacturer’s instructions. Cells were passed through a 40 µm FLOWMI cell strainer (SP Bel-Art) and live (DAPI-negative), single cells sorted using FACS Aria II. Barcoding, RNA extraction and library preparation was carried out using Chromium Single Cell 3’ Reagent kits v3 (10X Genomics) according to the manufacturer’s instructions. Libraries were sequenced using an Illumina NextSeq550 with a mid-output Flowcell, 150 cycles (130M pair end reads, 2x 75bp). The read alignment was performed with Kallisto(61) using locally built index and transcript-to-gene mapping based on the GRCm38 (release 98) gene and genome assembly files from Ensembl. The unfiltered count matrices produced by Kallisto were converted to SingleCellExperiment (SCE) objects and empty droplets were removed with EmptyDrops(62) using the default parameters. After removing all genes with counts of zero for all barcodes, the remaining genes were annotated with relevant features from Ensembl. Cell-level QC was performed with Scater(63), excluding cells with high amount of reads coming from mitochondrial genes (nmads >3) (**Supplementary Fig. 8**). Using Seurat(64) we further filtered the count matrices to contain only protein-coding genes with non-zero counts in at least 10 cells and cells expressing at least 500 protein-coding genes, after which we merged the wild type and mutant libraries for each experiment, resulting in 2042 wild type and 2153 mutant cells for *Rpgr*^*ORF d*5^ and 2299 wild type and 3277 mutant cells for *Rpgr*^*Ex*3*d*8^ (**Supplementary Table 2**). Clustering of the cells for the two experiments was done with Seurat, by performing log-normalisation, centring and scaling, clustering based on the identified variable features using shared nearest neighbour graph and the Louvain algorithm with multilevel refinement with resolution = 0.5 to produce the UMAP plots (**Fig 1a and b**). Cluster annotation was initiated by identification of each cluster markers against the remaining clusters using Seurat’s FindMarkers function (with min.pct = 0.25, logfc.threshold = 0.25) and selecting only upregulated genes with *pvaladj* ≤ 0.05, followed by manual curation based on existing literature. The trajectory plots presented in Fig 1a were generated using Slingshot(65). Gene ontology analysis was performed using Enrichr(66–68).

### Immunohistochemistry

Mice were sacrificed, eyes enucleated and placed into Davidson’s fixative (28.5% ethanol, 2.2% neutral buffered formalin, 11% glacial acetic acid) for 1 hr (cryosectioning) or overnight (wax embedding). For cryosectioning, eyes were removed from Davidson’s fixative and placed into 20% sucrose in PBS buffer overnight for cryopreservation. Eyes were then embedded using optimal cutting temperature and kept at -80 °C until sectioned. For wax preservation, eyes were fixed in Davidson’s fixative overnight at 4 °C. Following fixation, eyes were incubated successively in 70% v/v, 80% v/v, 90% v/v and 100% v/v ethanol, twice in xylene and then paraffin, each for 45 min per stage, using a vacuum infiltration processor. After sectioning, paraffin was removed by washing with 100% xylene and sections were rehydrated through an ethanol series (100%, 80%, 50%, 30%, 5 min each). For antigen retrieval, sections were boiled in sodium citrate pH6.4 for 2 × 10 min, then cooled to room temperature and washed in dH2O. Sections were post-fixed in acetone for 10 min at -20 oC, then blocked/permeabilised with 4% BSA (Sigma) and 0.2% Triton-X100 (Fisher) for 1 hr at RT and incubated with primary antibodies overnight at 4 °C. Sections were then washed in PBST (PBS + 0.2% Triton-X100), incubated with secondary antibodies for 2 hr at RT, washed in PBST, incubated in DAPI for 5 min at RT and mounted with coverslips using Prolong Gold (Invitrogen). Primary antibodies used were: mouse anti-Rhodopsin (Abcam ab98887, 1:1000), GFAP (Abcam ab7260, 1:500), rabbit anti-p62 (ENZO BML-PW9860-0100, paraffin only, 1:300), rabbit anti-LAMP1 (Abcam ab24170, 1:500), rabbit anti-TMEM119 (Synaptic Systems 400 002, 1:500), rat anti-F4/80 (AbD Serotec MCA497EL, 1:500), rabbit anti-VDAC1 (Abcam ab15895, 1:200), rabbit anti-pMLKL (Abcam ab196436, 1:100), rabbit anti-cleaved caspase-3 (Cell Signaling Technology 9661, 1:400). Secondary antibodies used: Donkey anti-rabbit Alexa-488, Donkey anti-rabbit Alexa-594, Donkey anti-mouse Alexa-594, Donkey anti-rat Alexa-488 (all Invitrogen). Confocal imaging was performed on a Nikon A1+ Eclipse TiE inverted microscope or a Leica Stellaris inverted microscope. 40x dry and 60x oil immersion lenses were used. Data was acquired using NIS Elements AR software (Nikon Instruments Europe, Netherlands) or LASX software (Leica Biosystems). Z-stacks were processed and analysed in Fiji (ImageJ).

### Protein Extraction and Western Blotting

Retinas were lysed in 50 mM Tris pH8.0, 150 mM NaCl, 1% NP-40 buffer containing protease inhibitors and phosphatase inhibitors. Protein samples were separated by SDS-PAGE and electroblotted onto nitrocellulose membranes (Biorad) using the TransBlot Turbo System for 10 min (Biorad). Non-specific binding sites were blocked by incubation of the membrane with 5% non-fat milk in TBS containing 0.1% Tween 20 (TBST) for 1 hr. Proteins were detected using primary antibodies diluted in blocking solution (4% Bovine Serum Albumin in TBST). Primary antibodies used were: rabbit anti-AKT, rabbit anti-pAKT, rabbit anti-caspase 8, rabbit anti-cleaved-caspase 8 (all Cell Signalling Technology, 1:1000). Following washing in PBST, blots were incubated with the appropriate secondary antibodies conjugated to horseradish peroxidase (Pierce) and chemiluminescence detection of Super Signal West Pico detection reagent (Pierce) by high resolution image capture using the ImageQuant LAS4000 camera system (GE Healthcare). Images were transferred to Fiji/ImageJ and mean pixel intensity of protein bands measured for quantification, with an equal area of blot assessed across all bands.

### Transmission Electron Microscopy

6-, 12and 18-monthold mice were euthanised by transcardial perfusion using fixative (0.2M sodium cacodylate pH 7.4, 5% glutaraldehyde, 4% paraformaldehyde). Eyecups were enucleated and placed in 1 mL of fixative. After 30 min, the cornea and lens were removed, then left to incubate in fixative for a total of 2 hr at room temperature. Thereafter, retinas were either: 1. Washed in 0.1 M phosphate buffer (pH 7.4), post-fixed with 1% osmium tetroxide (Electron Microscopy Science) and dehydrated in an ethanol series prior to embedding in Medium Epoxy Resin (TAAB). Ultrathin (75 nm) sections of the retina were then stained with aqueous uranyl-acetate and lead citrate and then examined with a Hitachi 7000 electron microscope (Electron Microscope research services, Newcastle University Medical School). 2. Polymerised in 4% agarose (Genemate E-3126-25). 150 µm sections were collected into MilliQ water using a vibratome. Sections were then stained in 1% osmium tetroxide 0.1M sodium cacodylate for 40 min, with rocking at room temperature, covered. After rinsing in MilliQ water, the sections were stained with 1% uranyl acetate in 0.2 M maleate buffer, pH 6.0, for 1 hr, with rocking at room temperature, covered. The sections were rinsed in MilliQ water and dehydrated in a series of ethanol washes (50%, 70%, 90%, 100%, 100%) for 15 min each, followed by two 100% acetone washes, 15 min each. The sections were then embedded in Epon-12 resin by sandwiching the sections between two sheets of ACLAR (EMS 50425-10) and leaving them at 60 °C for 48 hr. Ultrathin silver sections (60 nm) were placed on copper slot grids (EMS FF2010-CU) and post-stained in 1.2% uranyl acetate in MilliQ water for 6 min, followed by staining in Sato’s lead (a solution of 1% lead acetate, 1% lead nitrate, and 1% lead citrate; all from Electron Microscopy Sciences) for 2 minutes. Sections were imaged on a JEOL JEM-1400 electron microscope.

### Mass Spectrometry

Dark-adapted mice were maintained in constant darkness for 12 hr overnight prior to retina harvesting (also performed in the dark with infrared illumination). Retinas were lysed in 2% SDS and kept at -80°C until testing. 2 retinas from 1 mouse were used per biological replicate, with 5 biological replicates per group (mutant and wild type). In all cases, cell lysate samples were prepared and analysed as described 23.

### Mitochondrial Stress Test Assay

Xfe24 Seahorse cartridges (Agilent) were hydrated overnight prior to the experiments in Seahorse XF Calibrant (Agilent) at 37 oC in a non-CO2 incubator. Mesh inserts from Xfe24 Islet plates were precoated in Cell-Tak (Corning) using the sodium bicarbonate adsorption method. On the day of the experiment, mice were sacrificed by dislocation of the neck, eyes were enucleated and the retinas dissected in PBS. 4-6 punches were cut from each retina using a 1 mm biopsy punch, transferred to precoated mesh inserts (ganglion cell side down) and placed in individual wells of Xfe24 Islet plates with 500 µl of Seahorse DMEM media pH 7.4 (Agilent) supplemented with 10 mM glucose, 2 mM pyruvate and 2 mM Glutamine. Biopsies were incubated at 37 oC in a non-CO2 incubator for 20 min prior to running the assay. 8-10 biopsies from 1 mouse were used as technical replicates for each experimental sample, with a total of 4 biological replicates (4 mutant and 4 wild type mice). Mitochondrial oxygen consumption rates were measured using an Agilent Seahorse Xfe/XF Analyser using the standard protocol (Mix, 2 min; Wait, 2 min; Measure, 3 min). After three basal OCR measurements were obtained, OCR was measured following sequential injection of mitochondrial drugs (3 measurements per drug injection); 2 µM oligomycin (port A), 1 µM FCCP (port B), 0.5 µM antimycin A + 0.5 µM rotenone (port C).

### Image analysis and statistics

Image analysis was performed using Fiji/ ImageJ. For punctate staining (p62, LAMP1, VDAC1) the standard deviation (sd) of pixel intensity was measured from sum intensity projections of the relevant channel within a defined region of interest (photoreceptor inner segments) to distinguish bright puncta from background autofluorescence. All sd measurements were normalised to mean pixel intensity in the DAPI channel within a defined area of the ONL. To quantify pMLKL positive cells, whole retina sections were imaged at 40x using the LASX Navigator function and cells were counted manually from maximum intensity projections of 6 µm z-stacks. Due to extreme thinning of the ONL in *Pde*6*b* mutants, cell counts were normalised to the total area of the ONL. All statistical analysis was carried out using GraphPad Prism 9 (version 9.5.1; GraphPad software, USA) as described in the text. To determine statistical significance, unpaired t-tests were used to compare between two groups, unless otherwise indicated. The mean ± the standard error of the mean (SEM) is reported in the corresponding figures as indicated. Statistical significance was set at P <0.05.

## Funding

We are grateful for support from Fight for Sight (FN: 5179/5180), the MRC (MH, LN, LM, PM: MCUU00007/14, MR/Y015002/1), the European Research Council (PM: grant agreement n°866355) and the Wellcome Trust (RM: 219607/Z/19/Z)

## Declaration of interests

The authors declare no competing interests.

## Supplementary Information

**Supplementary figure 1:**
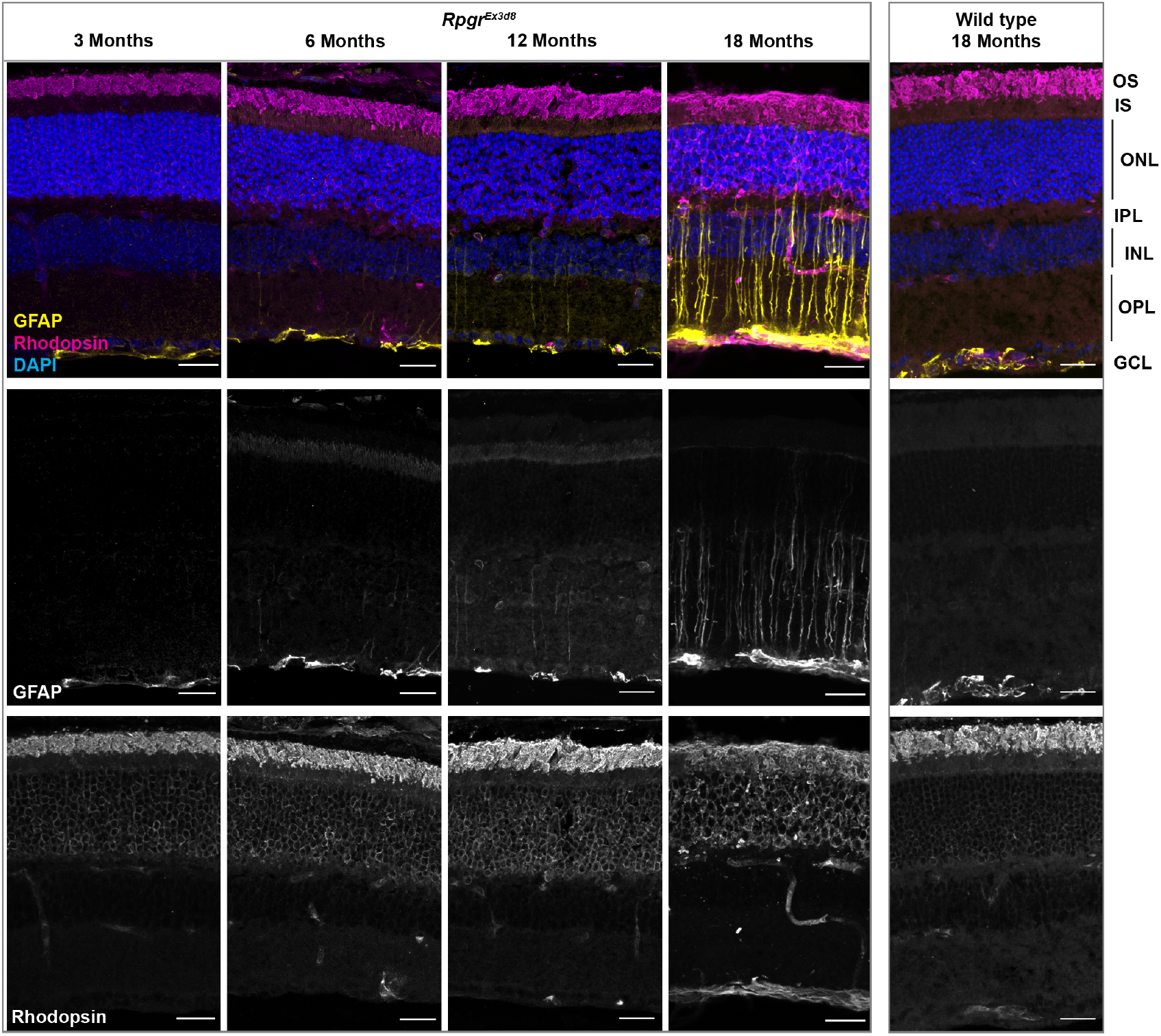
Rpgr mutants exhibit progressive retinal stress and rhodopsin mis-localisation. Reactive gliosis, as evidenced by increased GFAP immunolabelling throughout the radial length of Müller cells in the outer nuclear layer, increases in *Rpgr*^*Ex*3*d*8^ mutant retinas over time from 6 months of age (prior to significant ONL thinning at 18 months). Rhodopsin accumulates in inner segments and cell bodies at later time points (scale bars = 25 µm). OS = photoreceptor outer segments, IS = photoreceptor inner segments, ONL = outer nuclear layer, OPL = Outer plexiform layer, INL = inner nuclear layer, IPL = inner plexiform layer, GCL = ganglion cell layer.

**Supplementary figure 2:**
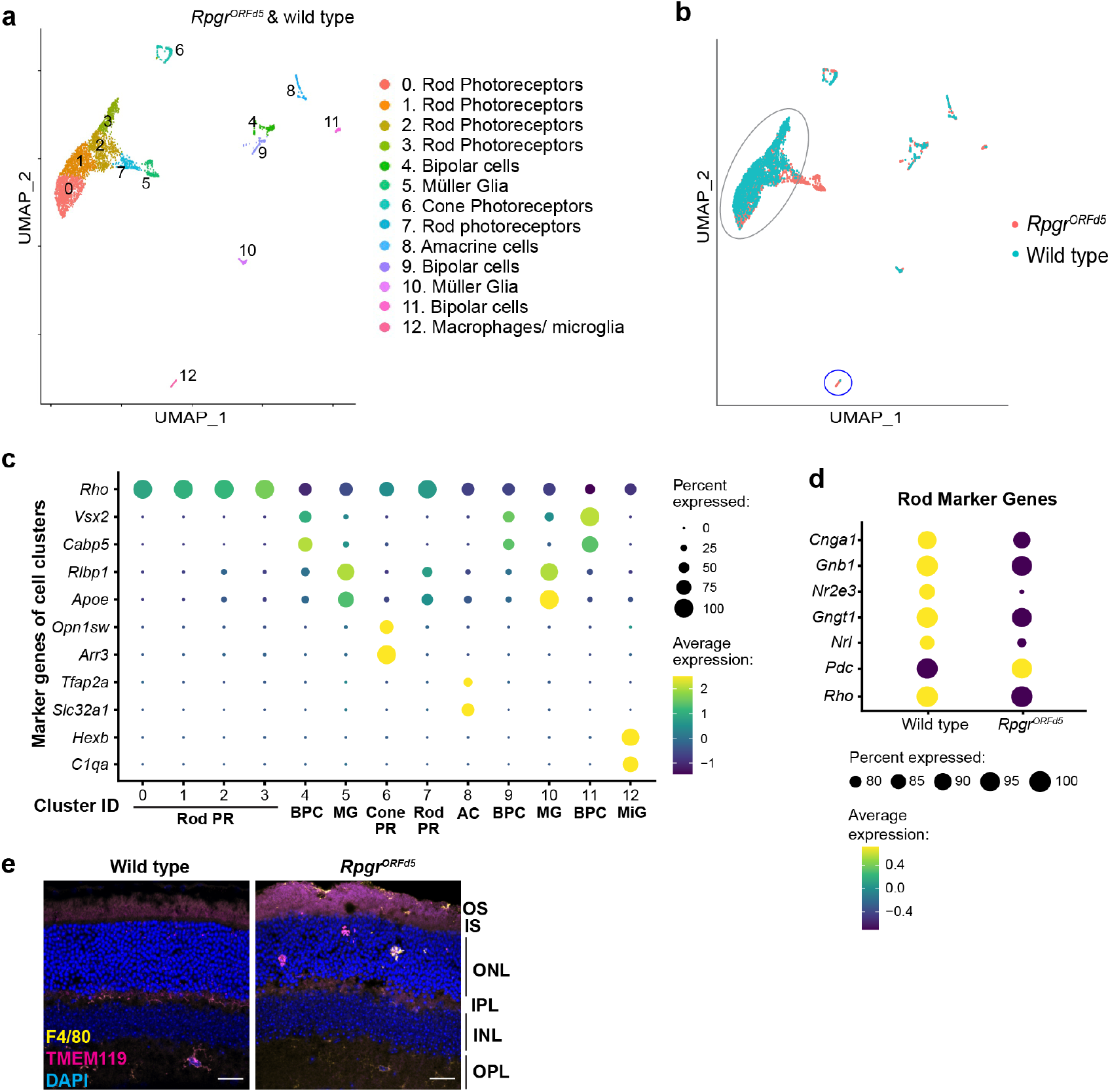
Single-cell transcriptomics identifies novel cell populations in *Rpgr*^*ORF d*5^ mutant retina. **a** UMAP plot showing clusters corresponding to photoreceptors and other retinal cell types in *Rpgr*^*ORF d*5^ retinas at 18 months (combined data from mutants and wild-type littermate controls, coloured by cluster identity, numbered by cluster size). **b** *Rpgr*^*ORF d*5^ mutant cells in salmon overlaid with wild-type cells in cyan. Rod photoreceptor clusters are circled in grey, macrophages (mainly *Rpgr*^*ORF d*5^ cells) circled in blue. **c** Dot plot showing expression of retinal cell type specific marker genes in each cluster (combined mutant and wild-type data, indicating cell type identity of each cluster (PR = photoreceptors, BPC = bipolar cells, MG = Müller glia, AC = amacrine cells, MiG = microglia/ macrophages). **d** Rod photoreceptor marker genes are downregulated in RpgrORFd5 mutant rod PR cells compared to wild type. **e** Macrophage cells (F4/80+; TMEM119-) and microglia (F4/80+; TMEM119+) are present in outer retinal layers in *Rpgr*^*ORF d*5^ mutants but not wild-type littermate controls at 18 months (Scale bar = 25 µm). OS = photoreceptor outer segments, IS = photoreceptor inner segments, ONL = outer nuclear layer, OPL = Outer plexiform layer, INL = inner nuclear layer, IPL = inner plexiform layer.

**Supplementary figure 3:**
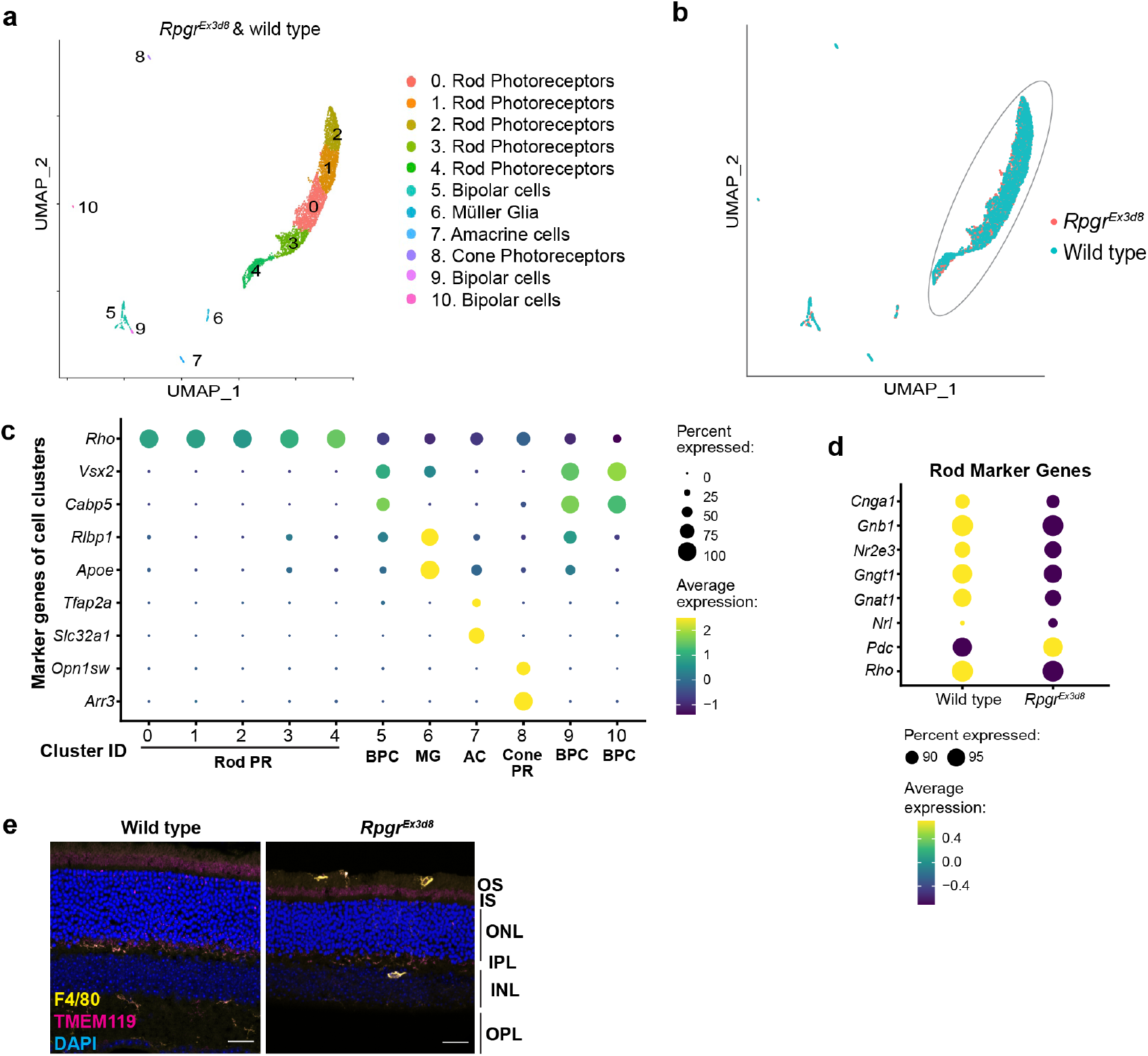
Single-cell transcriptomics identifies novel cell populations in *Rpgr*^*Ex*3*d*8^ mutant retina. **a** UMAP plot showing clusters corresponding to photoreceptors and other retinal cell types in *Rpgr*^*Ex*3*d*8^ retinas at 18 months (combined data from mutants and wild-type littermate controls, coloured by cluster identity, numbered by cluster size). **b** *Rpgr*^*Ex*3*d*8^ mutant cells in salmon overlaid with wild-type cells in cyan. Rod photoreceptor clusters are circled in grey. **c** Dot plot showing expression of retinal cell type specific marker genes in each cluster (combined mutant and wild-type data, indicating cell type identity of each cluster (PR = photoreceptors, BPC = bipolar cells, MG = Müller glia, AC = amacrine cells, MiG = microglia/ macrophages). **d** Rod photoreceptor marker genes are downregulated in RpgrORFd5 mutant rod PR cells compared to wild type. **e** Although macrophage cells were not present in the scRNAseq data for this mutant, macrophages (F4/80+; TMEM119-) and microglia (F4/80+; TMEM119+) are present in outer retinal layers in *Rpgr*^*Ex*3*d*8^ mutants but not wild-type littermate controls at 18 months (Scale bar = 25 µm). OS = photoreceptor outer segments, IS = photoreceptor inner segments, ONL = outer nuclear layer, OPL = Outer plexiform layer, INL = inner nuclear layer, IPL = inner plexiform layer.

**Supplementary figure 4:**
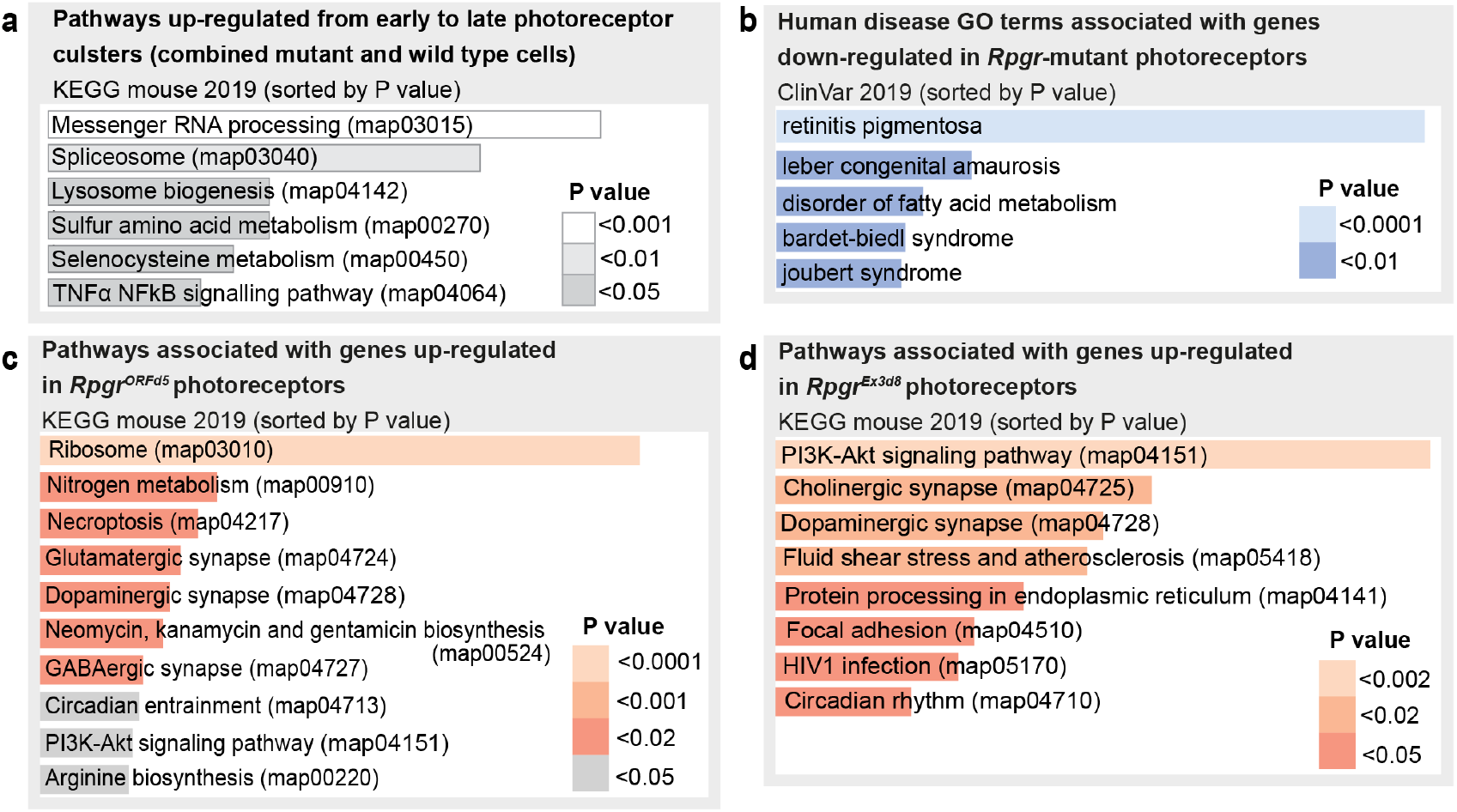
Cell stress pathways and PI3K-AKT signalling are up-regulated in degenerating and Rpgr mutant photoreceptors. GO term and KEGG pathway enrichment analyses performed using Enrichr 66–68. Bar length represents number of differentially expressed genes associated with each GO term/ KEGG pathway. **(a)** Pathways upregulated in rod photoreceptor subclusters along the disease trajectory: ‘early’ (less degenerated) to ‘late’ (more degenerated) in combined wild type and mutant cells. **(b)** Human disease GO terms associated with down-regulated genes identified from mutant photoreceptors compared to wild-type littermate control. **c, d** Pathways up-regulated in *Rpgr*^*ORF d*5^ **(c)** and *Rpgr*^*Ex*3*d*8^ **(d)** photoreceptors compared to wild-type littermate controls.

**Supplementary figure 5:**
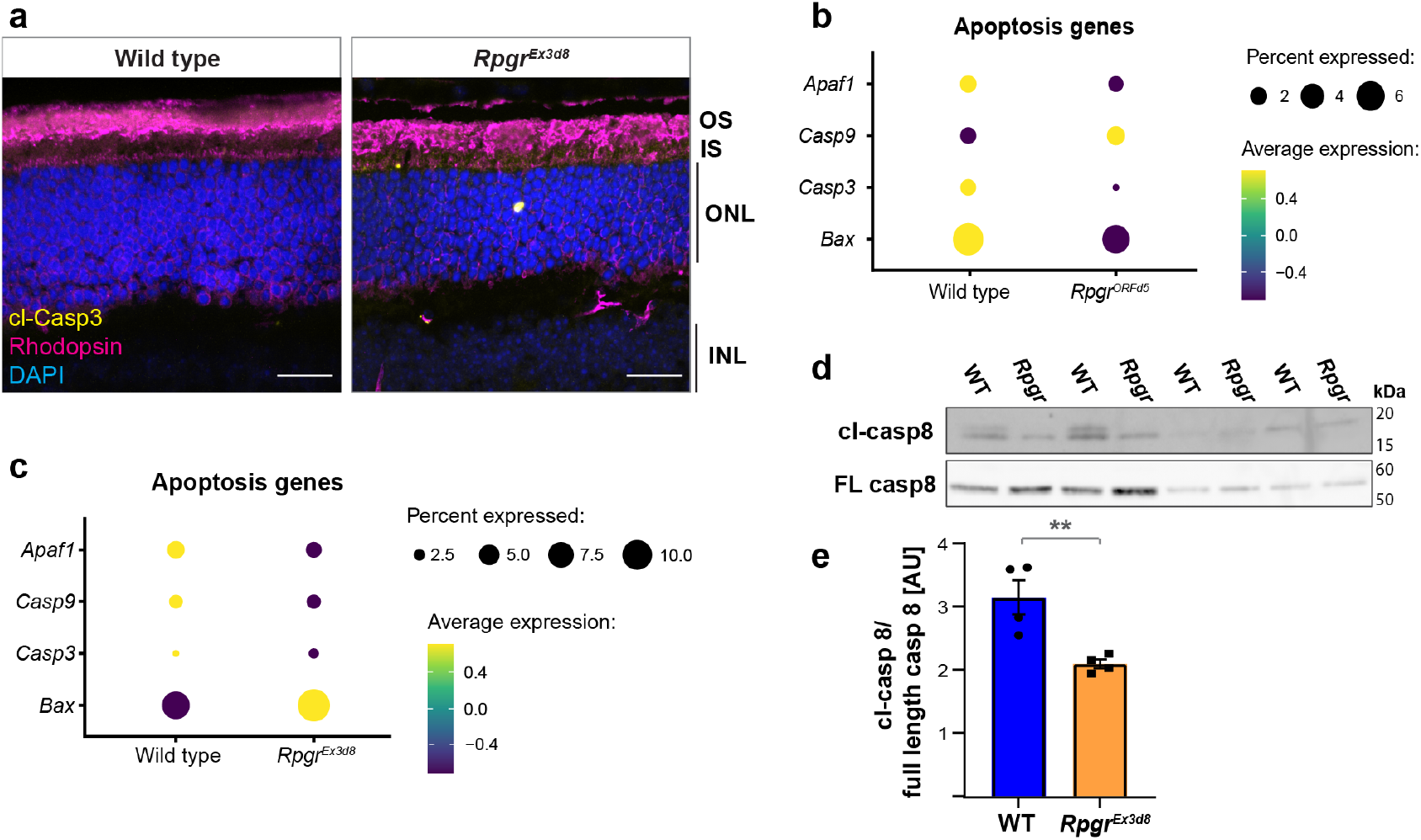
Apoptosis may not be the major cell death mechanism in Rpgr mutants. **a** Cleaved caspase-3 positive photoreceptors are present in *Rpgr*^*Ex*3*d*8^ mutant retinas at 18 months compared to wild type littermate controls. **b, c** Expression of apoptosis genes in *Rpgr*^*ORF d*5^ and *Rpgr*^*Ex*3*d*8^ mutants. **d, e** Immunoblot shows reduced cleaved caspase-8 relative to full-length caspase-8 in *Rpgr*^*Ex*3*d*8^ mutant retina lysates (N = 4 animals per experimental group; error bars show SEM; ** p = 0.0065).

**Supplementary figure 6:**
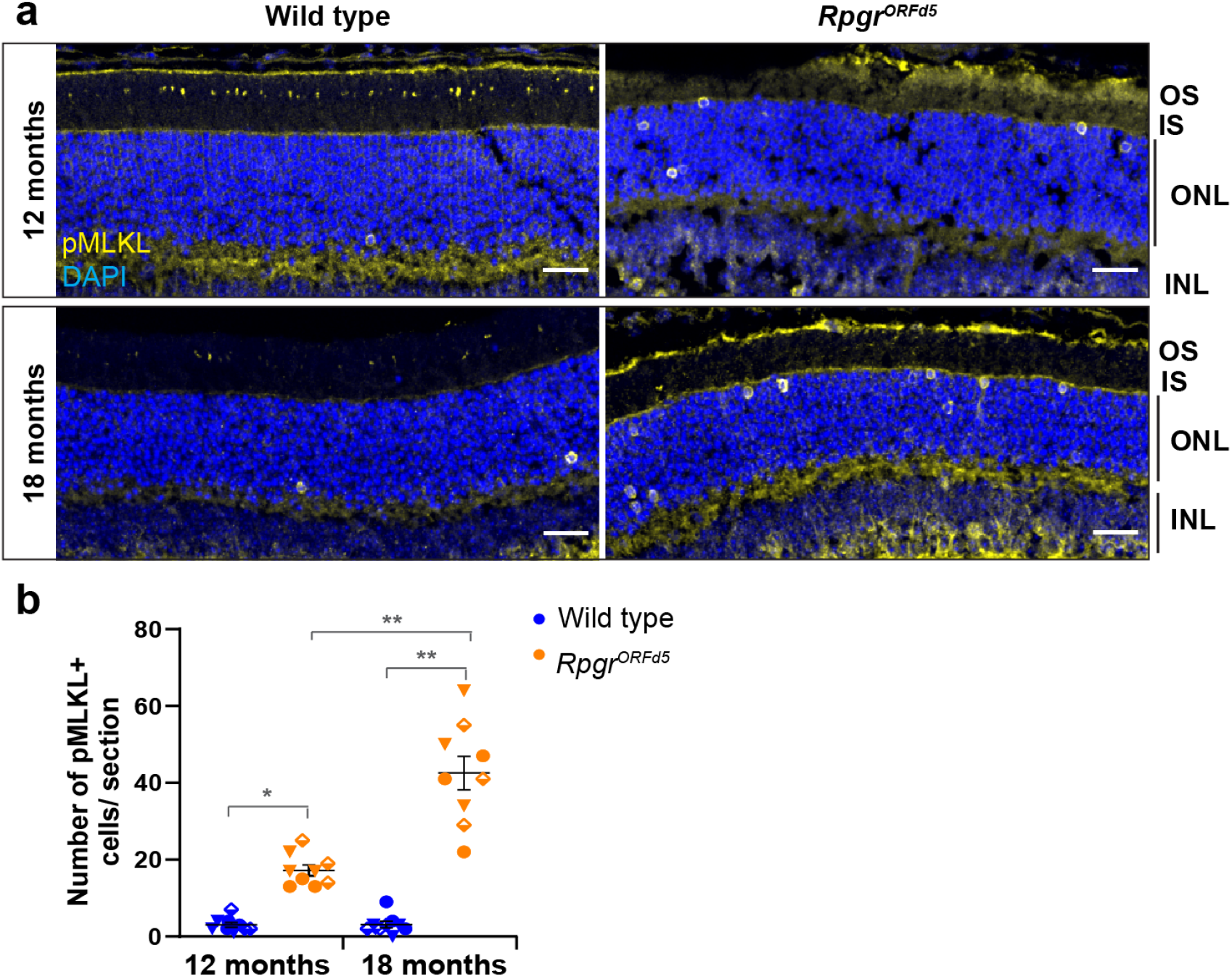
Progressive increase in photoreceptor necroptosis in *Rpgr*^*ORF d*5^ mutants. **a** Increased pMLKL positive photoreceptors in *Rpgr*^*ORF d*5^ mutants at 12 and 18 months compared to wild type littermate controls (scale bar = 25µm). **b** Symbols indicate images from individual mice (N = 3 animals per condition, >2 sections per animal; bars show mean; error bars show SEM; * p = 0.012; ** p = 0.008 (18 months), p = 0.009 (12 vs 18 months)).

**Supplementary figure 7.**
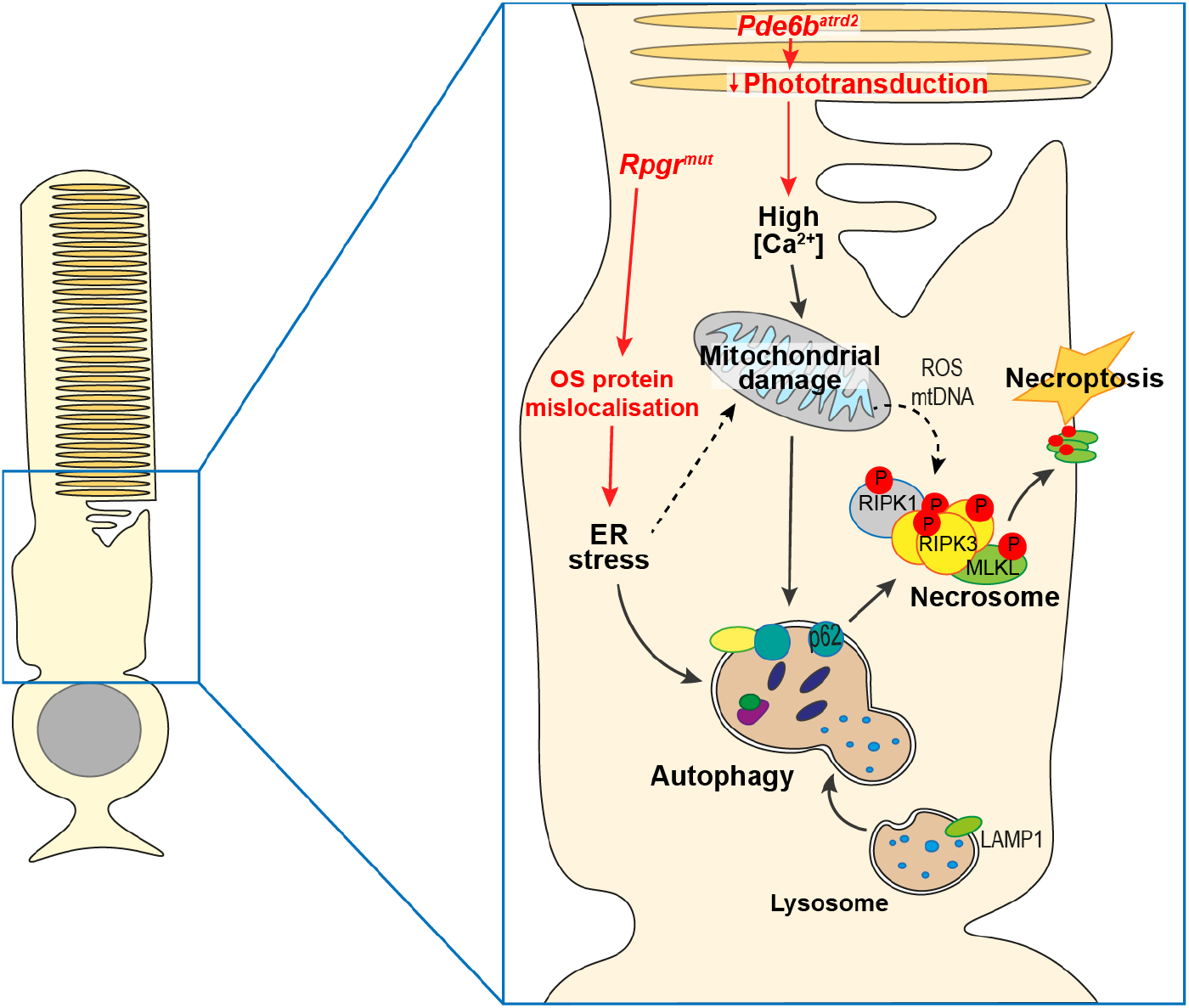
Different mechanisms of photoreceptor damage in *Rpgr* and *Pde*6*b*^*atrd*2^ mutants descend on a common cell death pathway. Phototransduction is impaired in *Pde*6*b*^*atrd*2^ mutants, possibly leading to raised intracellular calcium and mitochondrial damage. Outer segment maintenance is impaired in *Rpgr* mutants, leading to mislocalisation of OS proteins, increased endoplasmic reticulum (ER) stress and mitochondrial damage. Autophagy, to remove damaged mitochondria, increases in early disease. Necroptosis is facilitated by increased autophagy (via interaction of p62 with the necrosome), and release of reactive oxygen species (ROS) and mitochondrial DNA (mtDNA) from damaged mitochondria, facilitating necrosome formation.

**Supplementary figure 8:**
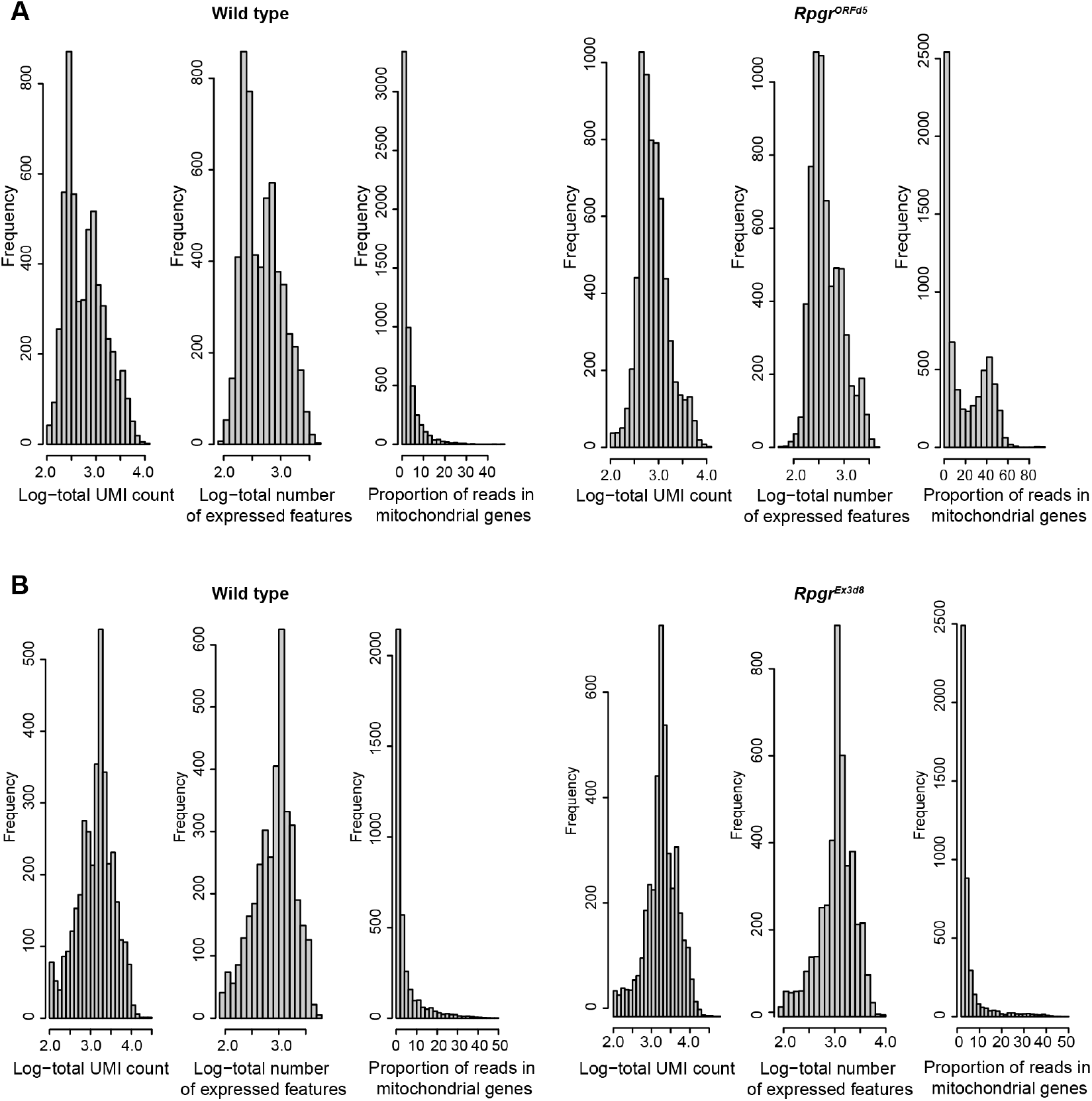
Cell-level quality control performed by Scater. Diagnostic plots for cell-level QC in scRNAseq data for *Rpgr*^*ORF d*5^ **a** and *Rpgr*^*Ex*3*d*8^ **b** retinas investigating the distribution of total number of counts for the cell (Log-total UMI count), the total number of features for the cell (Log-total number of expressed features) and the percentage of all counts for the cell that come from mitochondrial genes (Proportion of reads in mitochondrial genes).

**Supplementary Table 1:**
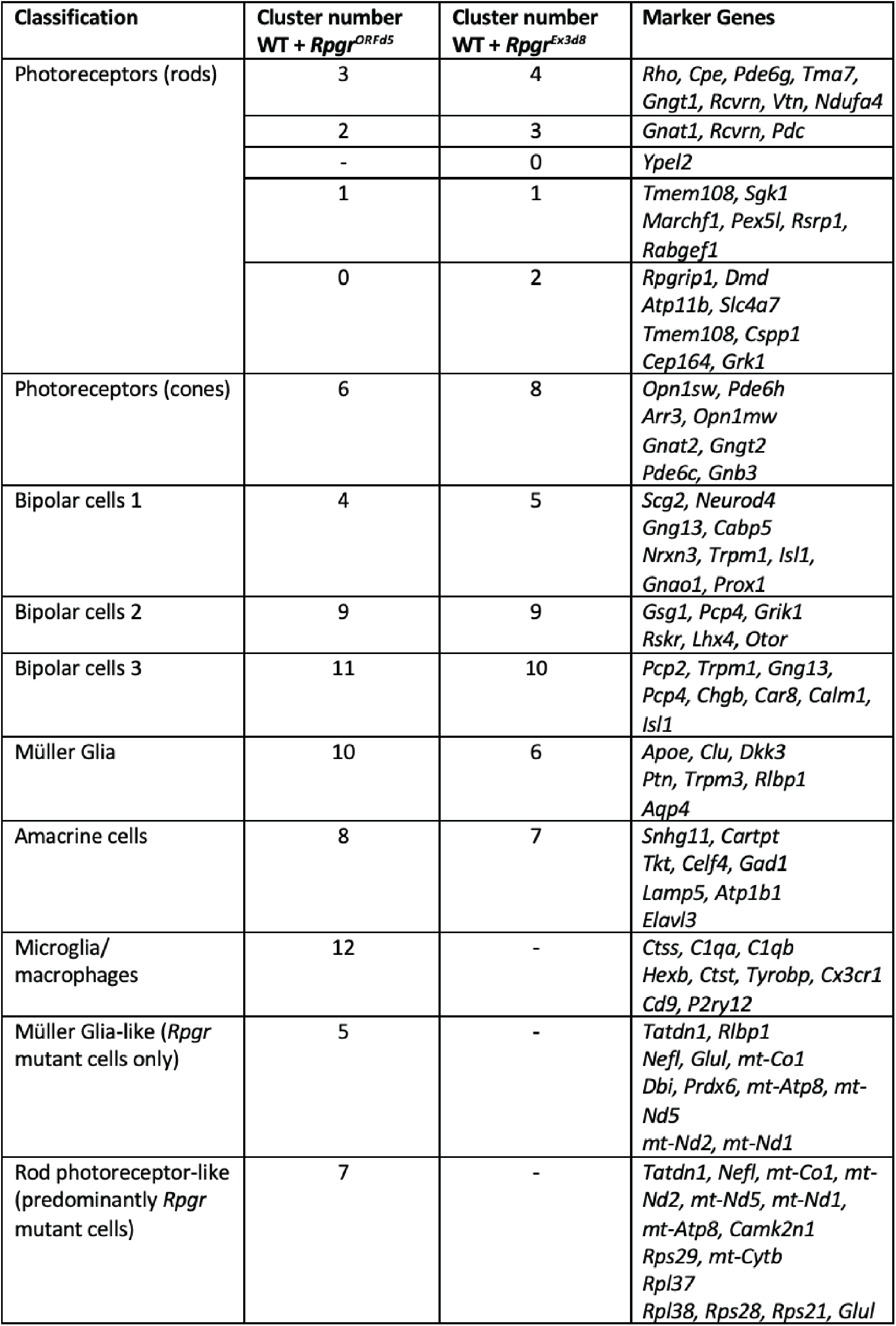
Marker genes used to identify each cluster as defined by Seurat. Genes listed are the most significantly expressed genes in each cluster compared to all other clusters (ranked by average log2FC, adjusted *pvalue* ≤ 0.0001) and significantly expressed in equivalent clusters in both experiments (if equivalent clusters were present).

**Supplementary Table 2A:** Number of WT and *Rpgr* mutant cells in each cluster as defined by Seurat.

| Cluster | WT<br>Cell Number | *Rpgr*<sup>ORFd5</sup> Mut<br>Cell Number | Cluster | WT<br>Cell Number | *Rpgr*<sup>Ex3d8</sup> Mut<br>Cell Number |
| --- | --- | --- | --- | --- | --- |
| 0 | 525 | 562 | 0 | 594 | 811 |
| 1 | 526 | 304 | 1 | 504 | 759 |
| 2 | 390 | 248 | 2 | 448 | 533 |
| 3 | 278 | 126 | 3 | 313 | 523 |
| 4 | 66 | 136 | 4 | 236 | 375 |
| 5 | 0 | 200 | 5 | 104 | 111 |
| 6 | 83 | 109 | 6 | 24 | 56 |
| 7 | 18 | 168 | 7 | 29 | 29 |
| 8 | 72 | 66 | 8 | 19 | 33 |
| 9 | 47 | 84 | 9 | 16 | 29 |
| 10 | 29 | 46 | 10 | 12 | 18 |
| 11 | 3 | 59 |  |  |  |
| 12 | 5 | 45 |  |  |  |
| **Total** | **2042** | **2153** |  | **2299** | **3277** |

**Supplementary Table 2B:** Percentage of *Rpgr* mutant and WT rod photoreceptors within each subcluster.

| Cluster | %<br>WT rod PR | %<br>*Rpgr*<sup>ORFd5</sup><br>rod PR | Cluster | %<br>WT rod PR | %<br>*Rpgr*<sup>Ex3d8</sup><br>rod PR |
| --- | --- | --- | --- | --- | --- |
| 0 | 30.5 | 45.3 | 0 | 28.4 | 27.0 |
| 1 | 30.6 | 24.5 | 1 | 24.1 | 25.3 |
| 2 | 22.7 | 20.0 | 2 | 21.4 | 17.4 |
| 3 | 16.2 | 10.2 | 3 | 14.9 | 17.4 |
|  |  |  | 4 | 11.3 | 12.5 |

**Supplementary Table 3:** Genes encoding mitochondrial proteins are up-regulated in *Rpgr* mutant rod photoreceptors. 6 out of 26 genes (23%) significantly up-regulated in *Rpgr*^*Ex*3*d*8^ mutant photoreceptors (adjusted p value < 0.0001) and 14 out of 63 genes (22%) significantly up-regulated in *Rpgr*^*ORF d*5^ mutant photoreceptors are involved in mitochondrial function.

| *Rpgr*<sup>ORFd5</sup> |  |  | *Rpgr*<sup>Ex3d8</sup> |  |  |
| --- | --- | --- | --- | --- | --- |
| Gene | log<sub>2</sub>FC | pVal adj | Gene | log<sub>2</sub>FC | pVal adj |
| *mt-Co1* | -1.4893338 | 3.73E-109 | *mt-Nd3* | -0.4191463 | 3.06E-19 |
| *mt-Atp8* | -1.3587884 | 1.50E-118 | *Bcl2* | -0.3311563 | 1.76E-22 |
| *mt-Nd2* | -1.2949704 | 4.02E-108 | *mt-Cytb* | -0.3260711 | 5.16E-11 |
| *mt-Nd5* | -1.2383093 | 1.88E-92 | *mt-Nd4* | -0.3015749 | 5.72E-10 |
| *mt-Nd1* | -1.1239659 | 4.90E-82 | *mt-Nd2* | -0.2583465 | 1.24E-06 |
| *mt-Nd4* | -1.0520625 | 3.74E-80 | *mt-Nd1* | -0.258555 | 4.79E-09 |
| *mt-Nd3* | -1.0380284 | 4.37E-69 |  |  |  |
| *mt-Nd6* | -0.9636443 | 3.41E-66 |  |  |  |
| *mt-Cytb* | -0.9427124 | 1.04E-68 |  |  |  |
| *mt-Nd4l* | -0.8190687 | 3.48E-49 |  |  |  |
| *Hk2* | -0.4155695 | 2.22E-18 |  |  |  |
| *Ppia* | -0.4082531 | 6.39E-14 |  |  |  |
| *Atp5k* | -0.3452921 | 9.81E-10 |  |  |  |
| *Vdac1* | -0.3452309 | 3.54E-07 |  |  |  |

**Supplementary Table 4:**
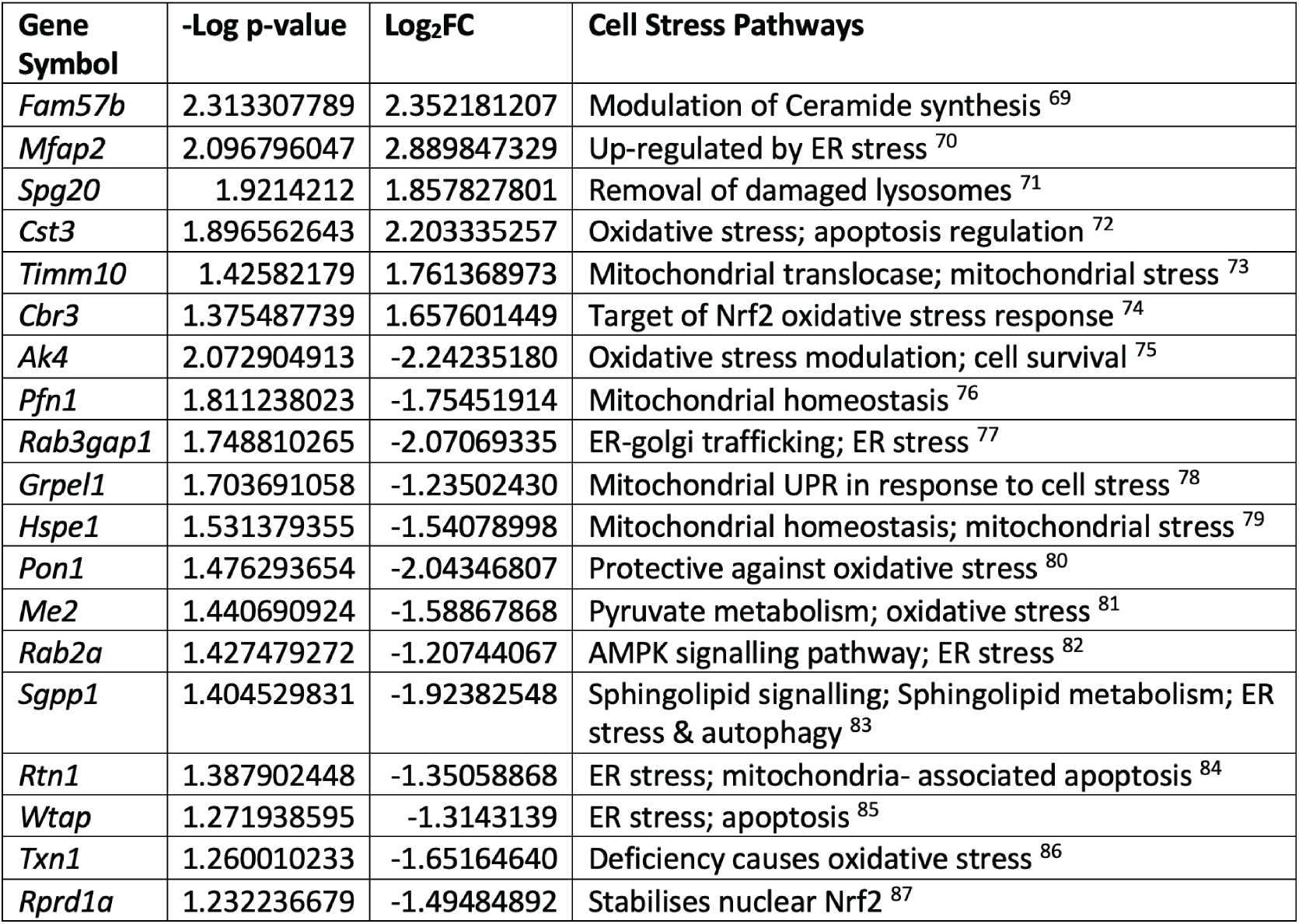
Differentially expressed proteins in *Rpgr*^*Ex*3*d*8^ retina compared to wild type have roles in cell stress pathways. Genes associated with differentially expressed peptides identified in whole retina mass spectrometry 23 (>1.5 fold enrichment (log2FC > 0.6 or <-0.6) and *p* ≤ 0.05) with functions in cell stress pathways. 6 out of 31 (19%) significantly up-regulated proteins and 13 out of 82 (16%) down-regulated proteins in *Rpgr*^*Ex*3*d*8^ retina are associated with cell stress pathways, particularly oxidative and mitochondrial stress.

| Gene Symbol | -Log p-value | Log<sub>2</sub>FC | Cell Stress Pathways |
| --- | --- | --- | --- |
| *Fam57b* | 2.313307789 | 2.352181207 | Modulation of Ceramide synthesis <sup>69</sup> |
| *Mfap2* | 2.096796047 | 2.889847329 | Up-regulated by ER stress <sup>70</sup> |
| *Spg20* | 1.9214212 | 1.857827801 | Removal of damaged lysosomes <sup>71</sup> |
| *Cst3* | 1.896562643 | 2.203335257 | Oxidative stress; apoptosis regulation <sup>72</sup> |
| *Timm10* | 1.42582179 | 1.761368973 | Mitochondrial translocase; mitochondrial stress <sup>73</sup> |
| *Cbr3* | 1.375487739 | 1.657601449 | Target of Nrf2 oxidative stress response <sup>74</sup> |
| *Ak4* | 2.072904913 | -2.24235180 | Oxidative stress modulation; cell survival <sup>75</sup> |
| *Pfn1* | 1.811238023 | -1.75451914 | Mitochondrial homeostasis <sup>76</sup> |
| *Rab3gap1* | 1.748810265 | -2.07069335 | ER-golgi trafficking; ER stress <sup>77</sup> |
| *Grpel1* | 1.703691058 | -1.23502430 | Mitochondrial UPR in response to cell stress <sup>78</sup> |
| *Hspe1* | 1.531379355 | -1.54078998 | Mitochondrial homeostasis; mitochondrial stress <sup>79</sup> |
| *Pon1* | 1.476293654 | -2.04346807 | Protective against oxidative stress <sup>80</sup> |
| *Me2* | 1.440690924 | -1.58867868 | Pyruvate metabolism; oxidative stress <sup>81</sup> |
| *Rab2a* | 1.427479272 | -1.20744067 | AMPK signalling pathway; ER stress <sup>82</sup> |
| *Sgpp1* | 1.404529831 | -1.92382548 | Sphingolipid signalling; Sphingolipid metabolism; ER stress & autophagy <sup>83</sup> |
| *Rtn1* | 1.387902448 | -1.35058868 | ER stress; mitochondria- associated apoptosis <sup>84</sup> |
| *Wtap* | 1.271938595 | -1.3143139 | ER stress; apoptosis <sup>85</sup> |
| *Txn1* | 1.260010233 | -1.65164640 | Deficiency causes oxidative stress <sup>86</sup> |
| *Rprd1a* | 1.232236679 | -1.49484892 | Stabilises nuclear Nrf2 <sup>87</sup> |

## Notes

### Competing Interest Statement

The authors have declared no competing interest.

